# *ADP-ribose-acceptor hydrolase 2* (*Arh2*) deficiency results in cardiac dysfunction, tumorigenesis, inflammation, and decreased survival

**DOI:** 10.1101/2023.02.07.527494

**Authors:** Jiro Kato, Sachiko Yamashita, Hiroko Ishiwata-Endo, Shunya Oka, Zu-Xi Yu, Chengyu Liu, Danielle A. Springer, Audrey Noguchi, Morteza Peiravi, Victoria Hoffmann, Martin J. Lizak, Matthew Medearis, In-Kwon Kim, Joel Moss

## Abstract

ADP-ribosylation is a reversible reaction with ADP-ribosyltransferases catalyzing the forward reaction and ADP-ribose-acceptor hydrolases (ARHs) hydrolyzing the ADP-ribose acceptor bond. ARH2 is a member of the 39-kDa ARH family (ARH1-3), which is expressed in heart and skeletal muscle. ARH2 failed to exhibit any in vitro enzymatic activity. To determine its possible in vivo activities, *Arh2*-knockout (KO) and - heterozygous (Het) mice were generated using CRISPR-Cas9. *Arh2*-KO mice exhibited decreased cardiac contractility by MRI, echocardiography and dobutamine stress with cardiomegaly and abnormal motor function. *Arh2*-Het mice showed results similar to those seen in *Arh2*-KO mice except for cardiomegaly. *Arh2*-KO and -Het mice and mouse embryonic fibroblasts (MEFs) developed spontaneous tumors and subcutaneous tumors in nude mice. We identified 13 mutations in *Arh2*-Het MEFs and heterozygous tumors, corresponding to human *ARH2* mutations in cancers obtained from COSMIC. Of interest, the L116R mutation in *Arh2* gene plays a critical role in aggressive tumorigenesis in nude mice, corresponding to human *ARH2* mutations in stomach adenocarcinoma. Both genders of *Arh2*-KO and -Het mice showed increased unexpectedly deaths and decreased survival rate during a 24-month observation, caused by tumor, inflammation, non-inflammation (e.g., cardiomegaly, dental dysplasia), and congenital diseases. Thus, *Arh2* plays a pivotal role in cardiac function, tumorigenesis, inflammation, and overall survival.

## Introduction

ADP-ribosylation is a post-translational modification where the ADP-ribose of NAD^+^ is transferred to an acceptor (e.g., protein) (1). Both mono- and poly-(ADP-ribosyl)ation involved in critical biological processes (2, 3). Arginine-specific mono-ADP-ribosyltransferases (ARTs) (e.g., ART1, ART5) catalyze mono-(ADP-ribosyl)ation (4, 5), forming an α-anomeric *N*-glycosidic linkage of ADP-ribose to arginine (6, 7). Poly(ADP-ribose) polymerases (PARPs) (e.g., PARP1, PARP2) catalyze poly-(ADP-ribosyl)ation, forming a branched, long-chain poly(ADP-ribose) polymer (PAR) (8, 9). ADP-ribosylation can be reversed by various hydrolase enzymes including poly-(ADP-ribose) glycohydrolase (PARG) (10), macrodomain-containing proteins (11, 12), and ADP-ribosyl-acceptor hydrolases (ARHs) (13, 14).

ARH2, also known as ADPRHL1, is a member of the 39-kDa ARH family (ARH1-3), is expressed mainly in heart and skeletal muscle (15–17). Although ARH1 and ARH3 catalyze enzymatic activities (ARH1 cleaves ADP-ribose-arginine, PAR, and *O*-acetyl-ADP-ribose (*O*AADPr); ARH3 cleaves PAR, ADP-ribose-serine, and *O*AADPr) in a Mg^2+^-dependent manner (18–24), to date ARH2 lacks any enzymatic activity (19, 25, 26). Consistently, ARH2 does not contain critical residues involving in Mg^2+^ binding in a sequence alignment (19).

Of interest, recent studies showed that ARH2 appears to be involved in myocardial function and tumorigenesis. Whole-genome sequencing analysis of 405,732 electrocardiograms from 81,192 Icelanders showed that an *ARH2* missense mutation (p.Leu294Arg in *ARH2*) is involved in left anterior fascicular block (LAFB) (27). Study of gene knockdown or knockout in *Xenopus laevis* showed that ARH2 is essential for heart chamber outgrowth and myofibril assembly in ventricle cardiomyocytes (17, 28). In the carcinogenesis, whole-exome sequencing and the Cancer Genome Atlas (TCGA) database of the families revealed that *ARH2* mutation participates in the development of prostate cancer and uveal melanoma (29–31). An *Arh2*-deficient mouse model has not been reported to date. To understand better the role of ARH2, we generated *Arh2*-knockout (KO) and -heterozygous (Het) mice using CRISPR/Cas9 technique and investigated their phenotypes.

In this study, we described cardiac phenotypes and carcinogenicity in *Arh2*-KO and -Het mice. Both genders of *Arh2*-KO mice showed decreased cardiac contractility including cardiomegaly with cardiac fibrosis assessed by MRI, dobutamine stress tests measured by echocardiography, and histopathology, and decreased exercise capacity, skeletal muscle contraction strength, and motor coordination assessed by treadmill, isometric torque, and rotarod tests. Similar to the results seen in *Arh2*-KO mice, *Arh2*-Het male mice showed decreased cardiac contractility with dobutamine infusion and skeletal muscle contraction. To investigate whether *Arh2* deficiency in mice is involved in tumor formation, we monitored tumor development until age 24-month. *Arh2*-KO and -Het mice developed spontaneous tumors. *Arh2*-Het mouse embryonic fibroblasts (MEFs) injected into nude mice formed tumor corresponding to mutations in the remaining allele. We identified 18 mutations in those *Arh2*-Het MEFs, subcutaneous tumor in nude mice injected *Arh2*-Het MEFs and heterozygous spontaneous-tumor tissues. Specifically, 13 mutations corresponded to human *ARH2* mutations in various cancers obtained from the COSMIC database. Of interest, the L116R mutation in *Arh2* gene was found in one of cultured *Arh2*-Het MEFs and subcutaneous tumors derived from *Arh2*-Het MEFs in nude mice. The L116R mutation was present in the most rapidly growing *Arh2*-Het MEFs, and tumors in nude mice. The identical L116R mutation was found in the Catalogue Of Somatic Mutations In Cancer (COSMIC) (https://cancer.sanger.ac.uk/cosmic/gene/analysis?ln=ADPRHL1) database in a human stomach adenocarcinoma. Furthermore, *Arh2*-KO and -Het showed an increased incidence of unexpected deaths that might be caused by inflammation (e.g., endocarditis, brain meningitis), non-inflammation (e.g., cardiac infarction, liver necrosis, cardiomegaly, dental dysplasia), and congenital disease (e.g., hydrocephalus, malocclusion) compared to WT counterparts, resulting decreased survival rate during 24-month of age. These results suggested that ARH2 plays an essential role for the prevention of cardiac dysfunction, tumorigenesis, inflammatory/non-inflammatory diseases (e.g., cardiomegaly, dental dysplasia) in mice.

## Results

### Both genders of *Arh2*-KO mice showed cardiomegaly with myocardial fibrosis in an age-dependent manner

To determine the role of *in vivo* ARH2 activities, we generated *Arh2*-wild-type (WT), -heterozygous (Het), and -knockout (KO) mice using CRISPR-Cas9 system (Supplemental Methods and Supplemental Figure 1). According to the Human Protein Atlas, ARH2 protein is mainly expressed in human heart muscle and skeletal muscle (https://www.proteinatlas.org/ENSG00000153531-ADPRHL1/tissue). ARH2 protein was identified in the WT mouse heart and skeletal muscle, but not other organs by Western blotting and Northern blotting analysis (Supplemental Figure 1 and 2). Representative histological section of 8-month-old of *Arh2-*WT, -Het and -KO male hearts is shown in Figure 1A and B. Histopathology by H&E stain and Masson’s trichrome stain showed heart enlargement and myocardial interstitial fibrosis in *Arh2* KO mice hearts (Figure 1A and B). The percentage of cardiac fibrotic area in both genders of *Arh2*-KO mice hearts (male, 10.5%; female, 4.4%) was significantly increased compared to WT (male, 2.6%; female, 2.3%) and *Arh2*-Het (male, 3.7%; female, 3.1%) mouse hearts (Figure 1C). The frequency of cardiac fibrosis in both genders of *Arh2*-KO mouse hearts (male, 22.9%; female, 7.1%) was significantly greater than that in WT (male, 1.4%; female,1.5%) or *Arh2*-Het (male, 4.3%; female, 1.9%) mice hearts (Figure 1D). Consistent with the observation of heart enlargement, the heart weight/body weight (HW/BW) ratio in *Arh2*-KO mice was significantly greater than the WT and *Arh2*-Het mice, regardless of gender (Figure 1E). These results suggested that *Arh2*-KO mice exhibited cardiomegaly with cardiac fibrosis, regardless of gender. In contrast, both genders of 3-4-month-old mice among genotypes were not observed with either cardiac fibrosis or cardiomegaly (data not shown). Heart weight/body weight (HW/BW) ratio in both genders of 3-4-month-old mice among *Arh2* genotypes was not significant different (Supplemental Figure 3H). Thus, *Arh2*-KO mice exhibited cardiomegaly with fibrosis in an age-dependent manner.

**Figure 1.**
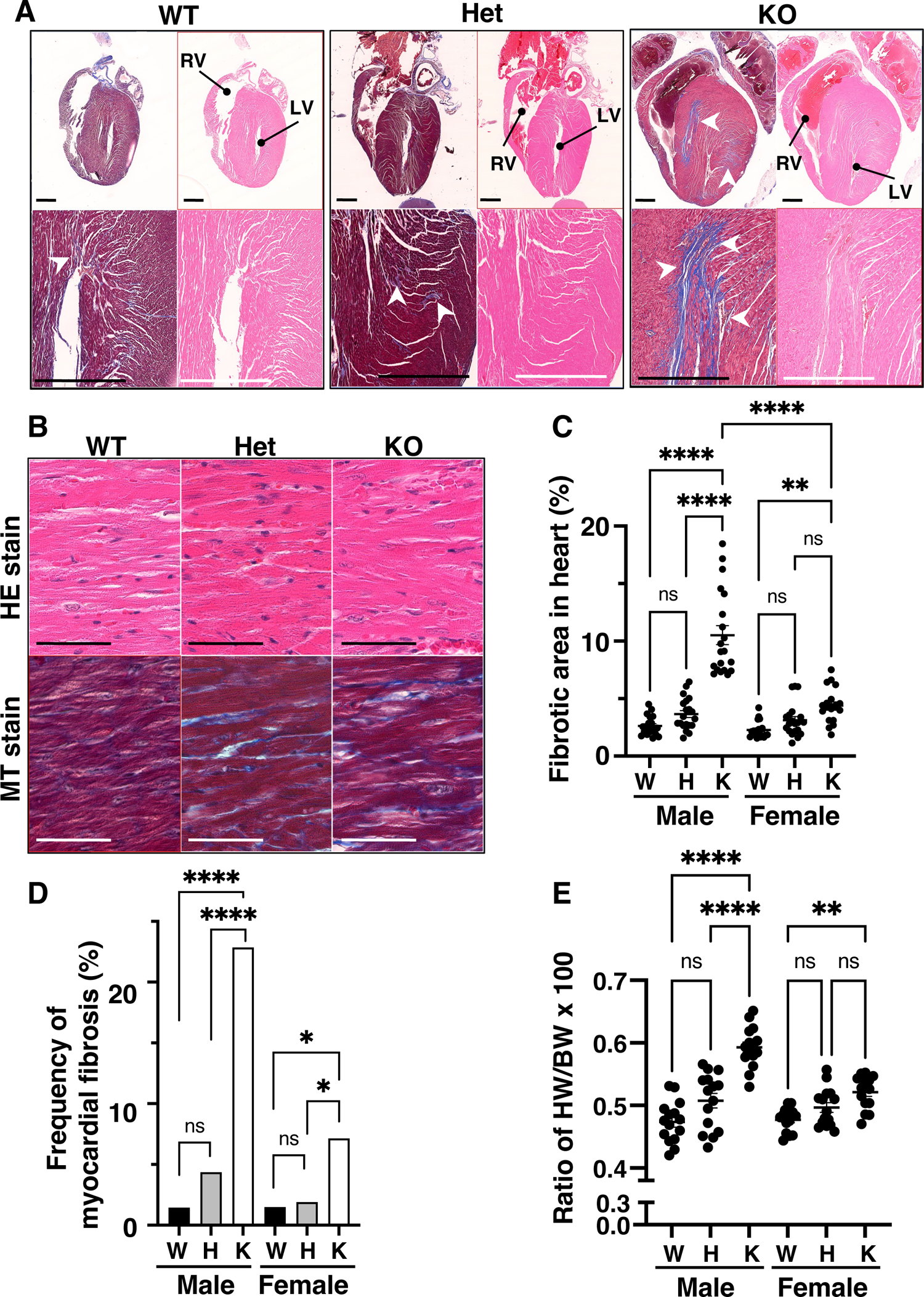
Cardiomegaly with myocardial fibrosis in *Arh2*-KO mouse heart. (**A**) Representative images of hematoxylin and eosin (HE) or Masson’s trichrome (MT) stained in heart sections of the 8-month-old, wild-type (WT), *Arh2*-heterozygous (Het), and *Arh2*-knockout (KO) male mouse. Left ventricular (LV) and right ventricular (RV) are indicated. Blue area in MT stained sections indicated fibrosis in heart (white arrowheads). Magnification of upper images and lower images in *Arh2* WT, Het, and KO male is 0.6x and 3.0x, respectively. Scale bars (black or white) show 1 mm. (**B**) High magnification images (40x) of HE and MT stained in heart sections of the 8-month-old, *Arh2* WT, Het and KO male mouse. Cardiac fibrosis was seen in *Arh2*-KO and -Het mouse heart. Scale bars (black or white) show 50 μm. (**C**) Percentage of fibrosis area in the whole cardiac chamber of MT stained sections was calculated by MetaMorpha software. Twenty individual heart sections from the 8-12-month-old mice among genotypes in both genders were collected. Wild-type (W), *Arh2*-heterozygous (H), and *Arh2*-knockout (K) mice are indicated. *n* = 20. Significant difference, ** *P* < 0.01, **** *P* < 0.0001 by one-way ANOVA, Tukey’s multiple comparison tests. (**D**) Frequency of myocardial fibrosis assessed by histopathological evaluation in WT (W, 1.5%, 2 of 138 males; 1.5%, 2 of 134 females), *Arh2*-Het (H, 4.3%, 7 of 161 males; 1.9%, 3 of 157 females), *Arh2*-KO (K, 22.9%, 16 of 70 males; 7.1%, 6 of 84 females). Significant difference **** *P* < 0.0001 by one-way ANOVA, Tukey’s multiple comparison tests. (**E**) Ratio of heart weight (HW) to body weight (BW) in the 7-9-months-old male and female mice among genotypes. *n* = 15. Significant difference, ** *P* < 0.01, **** *P* < 0.0001 by one-way ANOVA, Tukey’s multiple comparison tests.

**Figure 2.**
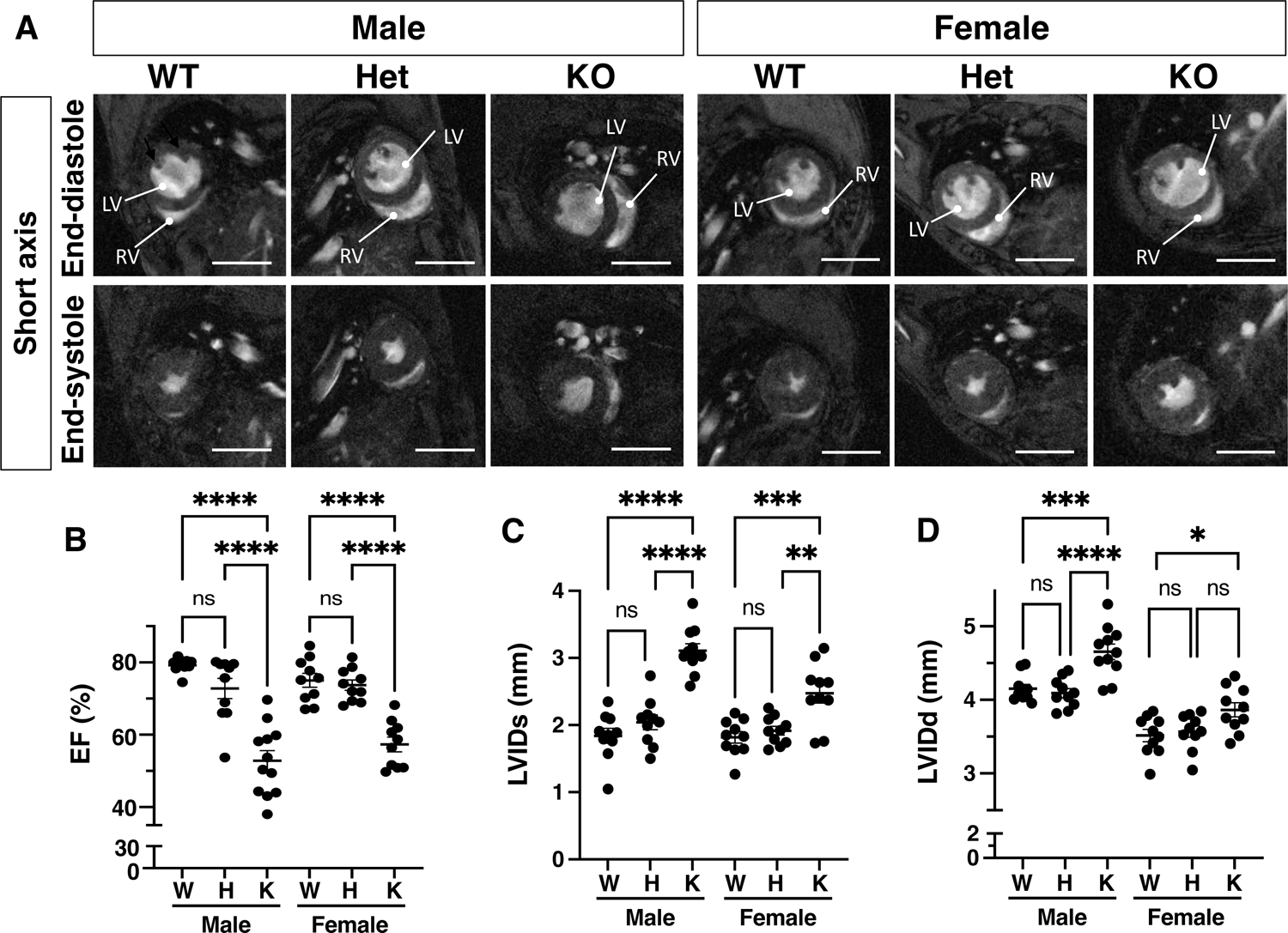
Both genders of *Arh2*-KO mice showed decreased cardiac contractility at baseline assessed by MRI. (**A**) Representative MRI images of short axis of mouse hearts at the end-systole and end-diastole in the 7-8-months-old male and female mice among *Arh2* genotypes. Left ventricle (LV) and right ventricle (RV) are indicated. Scale bars (5 mm, white). Corresponding MRI movies are shown in Supplemental video 1-6. (**B-D**) MRI measurements of cardiac contractility in the 7-8-months-old of male and female mice among *Arh2* genotypes. Wild-type (W), *Arh2*-heterozygous (H), and *Arh2*-knockout (K) mice are indicated. Cardiac MRI parameters: ejection fraction (EF, **B**), end-systolic left ventricle internal diameter (LVIDs, **C**), and end-systolic left ventricle internal diameter (LVIDs, **D**). *n* = 10-11. Significant difference, \**P* < 0.05, \*\**P* < 0.01, \*\*\**P* < 0.001, \*\*\*\**P* < 0.0001 by one-way ANOVA Tukey’s multiple comparison tests. ns: no significant.

### Both genders of Arh2-KO and -Het male mice showed decreased cardiac contractility in an age-dependent manner assessed by MRI and dobutamine stress by echocardiography

To determine the role of *Arh2* in myocardial function, cardiac contractility in the 7-8-month-old mice was evaluated by MRI and with a dobutamine stress test by echocardiography. Representative images of short axis of mouse hearts in systole and diastole, and video of cardiac contractility among genotypes in both male and female are shown in Figure 2A and online Supplemental Video 1. Based on the data analysis of MRI, the percentage of ejection fraction (%EF) in both genders of *Arh2*-KO mice were significantly decreased compared to those of WT and *Arh2*-Het mice (Figure 2B). Left ventricular internal diameter at end-systole (LVIDs) and LVID at diastole (LVIDd) in both genders of *Arh2*-KO mice were significantly greater than those of WT and *Arh2*-Het mice (Figure 2C and D). Thus, *Arh2*-KO mice showed decreased cardiac contractility regardless of gender at baseline assessed by MRI. In a dobutamine stress test monitored by echocardiography, *Arh2* mice were administered dobutamine (0, 10, and 40 μg/kg/min). Representative echocardiographic images of both genders of mouse hearts among genotypes at baseline are shown in Figure 3A. At baseline, %EF and % of fractional shortening (%FS) in both genders of *Arh2*-KO mice were significantly decreased compared to those in WT mice (Figure 3B and C). LVIDs, LVIDd, end-systolic volume (LVESV), and end-diastolic volume (LVEDV) in both genders of *Arh2*-KO mice were significantly greater than those in WT mice (Figure 3D-G). During pharmacological stress by dobutamine administration, cardiac contractility increased regardless of gender or genotypes in a dose-dependent manner (Figure 3D-G). Regardless of low- or high-dose dobutamine infusion, %EF and %FS in both genders of *Arh2*-KO mice were significantly decreased compared to those in WT and *Arh2*-Het mice (Figure 3B and C). LVIDs, LVIDd, LVESV, and LVEDV in both genders of *Arh2*-KO mice were significantly greater than those in WT mice (Figure 3D-G). In addition, %EF of *Arh2*-Het male mice was significantly decreased compared to those in WT mice during high-dose dobutamine infusion (Figure 3B). These results suggested that *Arh2*-KO mice showed decreased cardiac contractility with dobutamine stress regardless of gender. In addition, *Arh2*-Het male mice showed decreased cardiac contractility with dobutamine stress. In contrast, cardiac contractility (e.g., EF%, FS%) at baseline by echocardiography in 3-4-month-old of mice among genotypes was not significantly different (Supplemental Figure 3). Thus, both genders of *Arh2*-KO mice showed decreased cardiac contractility in an age-dependent manner.

**Figure 3.**
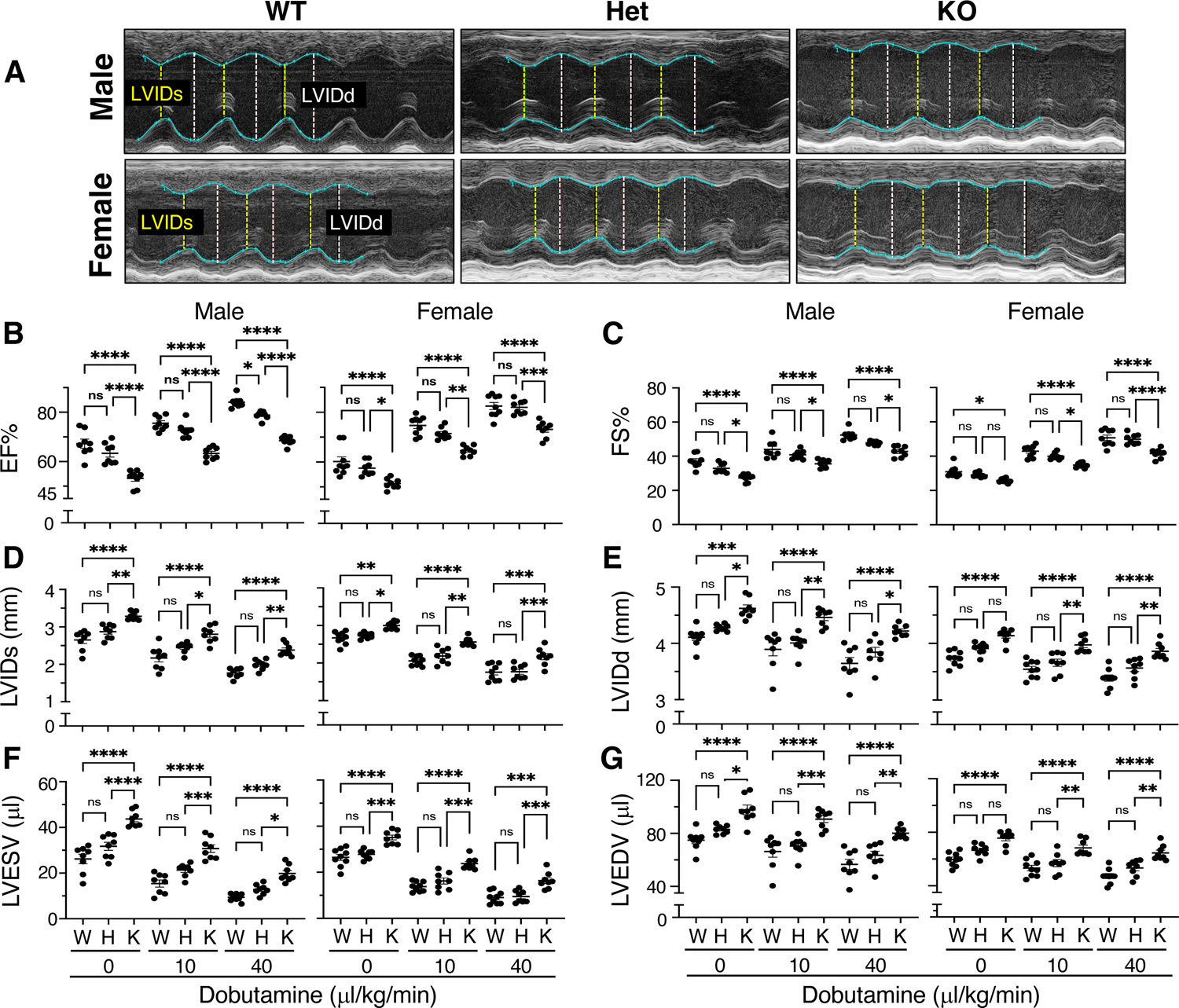
Both genders of *Arh2*-KO and -Het male mice showed decreased cardiac contractility assessed by echocardiography with dobutamine infusion. (**A**) Representative images by echocardiography at baseline in M-mode in the 7-8-months-old of male and female mice among *Arh2* genotypes. Wild-type (W), *Arh2*-heterozygous (H), and *Arh2*-knockout (K) mice are indicated. End-diastolic LVIDd, (white) and end-systolic LVIDs (yellow) are indicated. (**B-G**) Echocardiography analysis at dobutamine stress test (low-dose dobutamine: 10 μl/kg/min, high-dose dobutamine: 40 μl/kg/min) in the 7-8-months-old of male and female mice among *Arh2* genotypes. Cardiac echocardiography parameters: ejection fraction (%EF, **B**), fractional shortening (%FS, **C**), left ventricular internal diameter (mm) at systole (LVIDs, **D**), and at diastole (LVIDd, **E**), and left ventricular chamber volume (μL) at systole (LVESV, **F**) and at diastole (LVEDV, **G**). *n* = 8-9. Significant difference, \**P* < 0.05, \*\**P* < 0.01, \*\*\**P* < 0.001, \*\*\*\**P* < 0.0001 by one-way ANOVA Tukey’s multiple comparison tests. ns: no significant.

### Both genders of *Arh2*-KO and -Het male mice showed decreased exercise capacity, muscle contraction strength, and motor coordination assessed by treadmill, isometric torque, and rotarod tests

We showed that *Arh2*-KO mice displayed decreased cardiac contractility in both male and female in an age-dependent manner assessed by MRI and echocardiography. To determine the effect of ARH2 in muscle function, exercise capacity, muscle contraction strength, and motor coordination, 7-8-months-old *Arh2* mice were examined using treadmill, isometric torque, and rotarod tests. In the treadmill test, running time, total running distance, maximum speed, and capable of maximal work (kg/m) until exhaustion in both genders of *Arh2*-KO mice were significantly decreased compared to those in WT mice (Figure 4A-D). In addition, running time, total running distance, maximum speed, and capable of maximal work (kg/m) until exhaustion in *Arh2*-Het male mice were significantly decreased relative to those in WT mice (Figure 4A-D), but no difference was observed between female Het and WT mice. Thus, both genders of *Arh2*-KO mice and *Arh2*-Het male mice showed decreased exercise capacity. In the isometric torque test, specific torque in both genders of *Arh2-*KO mice was significantly decreased than that in *Arh2* WT during stimulation frequencies from 25 to 250 Hz (Figure 4E). In addition, specific torque in *Arh2*-Het male mice was significantly decreased than that in WT male mice (Figure 4E), but no difference was observed between female Het and WT mice. Thus, both genders of *Arh2*-KO mice and *Arh2*-Het male mice showed decreased muscle contraction strength. In the rotarod test, total run time in both genders of *Arh2*-KO mice was significantly reduced from that of WT, but no difference was observed between female Het and WT mice (Figure 4F). In addition, total run time in *Arh2*-Het male mice was significantly reduced from that of WT, but no difference was observed between female Het and WT mice (Figure 4F). Thus, both genders of *Arh2*-KO mice and *Arh2*-Het male mice showed decreased muscle coordination.

**Figure 4.**
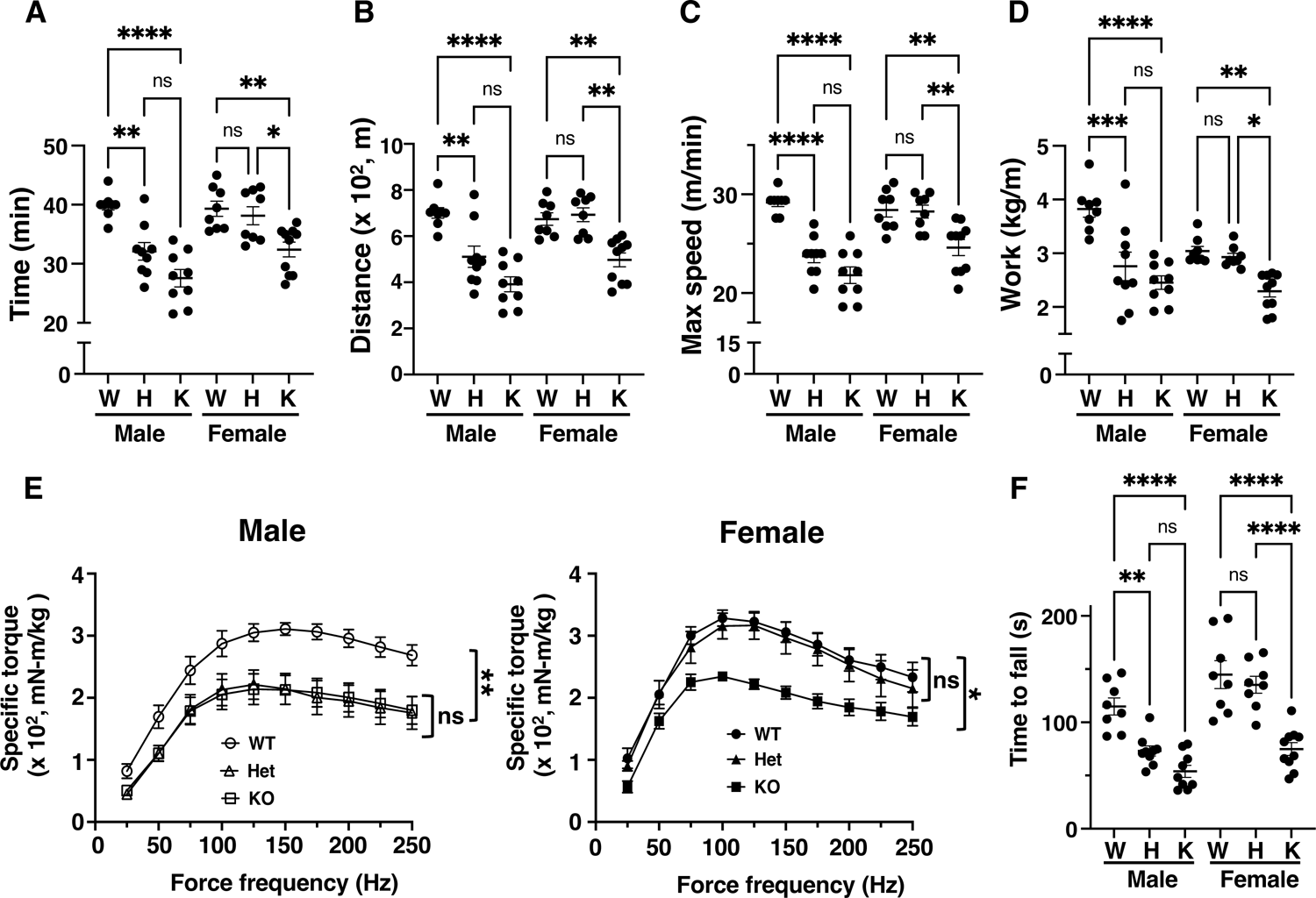
Both genders of *Arh2*-KO and -Het male mice showed decreased motor function assessed by treadmill, isometric torque, rotarod tests. (**A-D**) Exercise capacity assessed by treadmill test in the 7-8-months-old of male and female mice among *Arh2* genotypes. Wild-type (W), *Arh2*-heterozygous (H), and *Arh2*-knockout (K) mice are indicated. Measurements of running time (minutes, min) until exhaustion, (**A**); total running distance (meter, m), (**B**); maximum speed (meter per minutes, m/min), (**C**); work (kilogram per meter, kg/m) (**D**). *n* = 8-10. Significant difference, \**P* < 0.05, \*\**P* < 0.01, \*\*\**P* < 0.001, \*\*\*\**P* < 0.0001 by two-way ANOVA Tukey’s multiple comparison tests. ns: no significant. (**E**) Skeletal muscle contraction assessed by isometric torque test in the 7-8-months-old of male and female mice among *Arh2* genotypes. Wild-type (WT; ○, •), heterozygous (Het; Δ, Δ), and knockout (KO; □, ▀) are indicated. *n* = 8-9. Significant difference, \**P* < 0.05, \*\**P* < 0.01 by one-way ANOVA Tukey’s multiple comparison test. ns: no significant. (**F**) Motor coordination assessed by rotarod test in the 7-8-months-old of male and female mice among *Arh2* genotypes. *n* = 8-10. Significant difference, \**P* < 0.05, \*\**P* < 0.01 by one-way ANOVA Tukey’s multiple comparison test. ns: no significant.

### *Arh2*-KO and -Het mice result in a male predominant increase in spontaneous tumor, metastasis, and multiple tumors

To investigate tumor development, *Arh2* mice were monitored during a 24-month observation. Tumor classification was determined via histopathological evaluation. Tumors in *Arh2*-KO male mice appeared at the age of 1.5-month, increased until at 10-months when survival rate of *Arh2*-KO male mice reached to 0% due to unexpected deaths and inflammatory diseases (Figure 5A). Tumors in *Arh2*-Het male, -KO female, and -Het female mice appeared at the age of 5-months and increased until the 24-month observation (Figure 5A). Incidence of tumors in both genders of *Arh2*-KO (male, 42.9%; female, 29.8%) and Het mice (male, 39.1%; female, 19.1%) were significantly increased compared to WT mice (male, 4.3%; female, 3%) (Figure 5B). Of note, tumor incidence of *Arh2*-KO and Het male mice was significantly increased compared to female mice, suggesting that male is predominant in tumorigenesis (Figure 5B). A variety of spontaneous tumors (e.g., hematopoietic tumors-lymphoma, histiocytic sarcoma, hemangiosarcoma, leukemia; carcinoma-pulmonary adenocarcinoma, hepatocellular carcinoma) was observed in both genders of *Arh2* KO and Het mice, and only lymphoma was observed in WT mice (Figure 5B). Incidence of metastasis tumors in *Arh2-*KO mice (male, 21.4%; female, 10.7%) and *-*Het mice (male, 8.1%; female, 4.4%) was significantly increased than those in WT mice metastatic tumors (Figure 5C). Incidence of multiple tumors in *Arh2*-KO mice (male, 18.6%; female, 7.1%) and -Het mice (male, 8.1%; female, 2.5%) was significantly increased relative to those in WT mice (none seen) (Figure 5D). These results suggested that *Arh2*-KO and -Het mice showed a male predominant increase in spontaneous tumor, metastasis, and multiple tumors.

**Figure 5.**
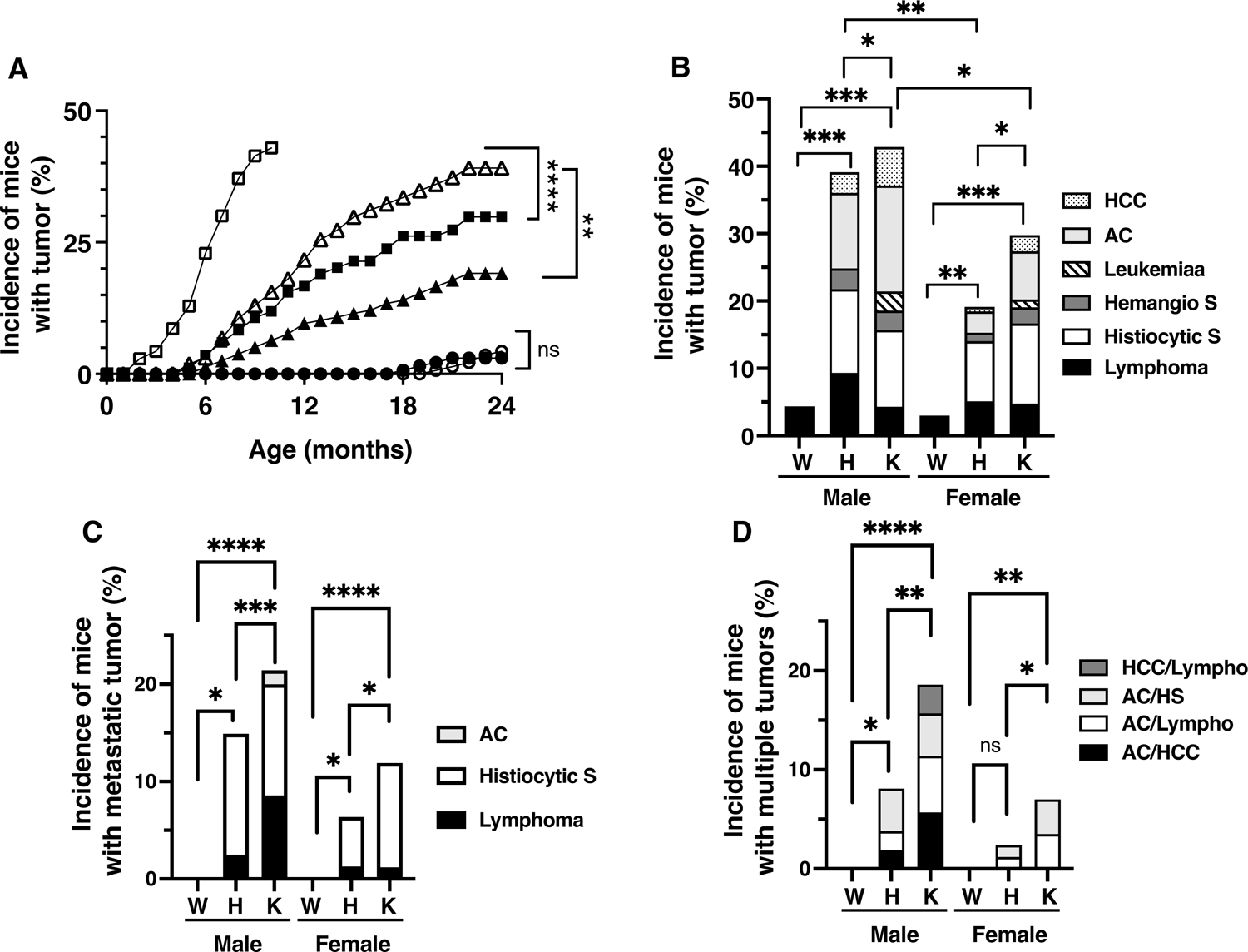
Both genders of *Arh2*-KO and -Het mice showed increased spontaneous tumor development. (**A**) Comparison the cumulative number of mice with tumor by *Arh2* genotypes and age during the 24-month observation. Tumors were subjected to histopathological evaluation. WT mice (male ○, female •), *Arh2*-Het mice (male Δ, female Δ), *Arh2*-KO mice (male □, female ▀) were indicated. The % of tumor incidence in WT mice (4.3%, 6 of 138 males; 3%, 4 of 134 females), *Arh2*-Het mice (39.1%, 63 of 161 males; 19.1%, 30 of 157 females), *Arh2*-KO mice (42.9%, 30 of 70 males; 29.8%, 25 of 84 females). Significant difference by one-way ANOVA Tukey’s multiple comparison test. Comparisons: *Arh2*-WT male mice vs *Arh2*-KO male mice, \*\*\*\**P* < 0.0001; *Arh2*-WT male mice vs *Arh2*-Het male mice, \*\*\*\**P* < 0.0001; *Arh2*-Het male mice vs *Arh2*-KO male mice, \*\*\**P* < 0.001; *Arh2*-WT female mice vs *Arh2*-KO female mice, \*\*\*\**P* < 0.0001; *Arh2*-WT female mice vs *Arh2*-Het female mice, \**P* < 0.05; *Arh2*-Het female mice vs *Arh2*-KO female mice, \**P* < 0.05; *Arh2*-KO male mice vs *Arh2*-KO female mice, \*\*\*\**P* < 0.0001; *Arh2*-Het male mice vs *Arh2*-Het female mice, \*\**P* < 0.01. ns; no significant between *Arh2*-WT male vs. WT-female. (**B**) Classification of tumor incidence in male and female mice among *Arh2* genotypes during the 24-month observation. Wild-type (W), *Arh2*-heterozygous (H), and *Arh2*-knockout (K) mice are indicated. lymphoma (black bar), histiocytic sarcoma (Histiocytic S, white bar), hemangiosarcoma (Hemangio S, dark-gray bar), leukemia (Leukemia, borders bar), adenocarcinoma (AC, light-gray bar), and hepatocellular carcinoma (HCC, dots bar) are indicated and % of tumor incidence is shown Supplemental Table 6. Significant difference, \**P* < 0.05, \*\**P* < 0.01, \*\*\**P* < 0.001, \*\*\*\**P* < 0.0001 by two-way ANOVA, Tukey’s multiple comparison tests. (**C**) Incidence of mice with metastasis tumor in male and female mice among *Arh2* genotypes during the 24-month observation. lymphoma (black bar), histiocytic sarcoma (Histiocytic S, white bar), and adenocarcinoma (AC, light-gray bar) are indicated. The % of metastasis tumor in *Arh2*-KO mice (21.4%, 15 of 70 males; 11.9%, 10 of 84 females) and *Arh2*-Het mice (14.9%, 24 of 161 males; 6.4%, 10 of 157 females). Metastasis tumor were not observed in WT male and female mice. Significant difference, \**P* < 0.05, \*\*\**P* < 0.001, \*\*\*\**P* < 0.0001 by two-way ANOVA, Tukey’s multiple comparison tests. (**D**) Incidence of mice with multiple tumors in male and female mice among *Arh2* genotypes during the 24-month observation. The % of multiple tumors in *Arh2*-KO mice (18.6%, 13 of 70 males; 7 %, 6 of 84 females) and *Arh2*-Het mice (8.1%, 13 of 161 males; 2.4%, 4 of 157 females) were observed. Multiple tumors were not observed in WT male and female mice. Adenocarcinoma and hepatocellular carcinoma (AC/HCC, black bar), adenocarcinoma and lymphoma (AC/Lympho, white bar), adenocarcinoma and histiocytic sarcoma (AC/HS, light-gray bar), and hepatocellular carcinoma and lymphoma (HCC/Lympho, dark-gray bar) are indicated. Significant difference, \**P* < 0.05, \*\**P* < 0.01, \*\*\*\**P* < 0.0001 by one-way ANOVA, Tukey’s multiple comparison tests. ns; no significant. Data of **A**-**D** are shown values from 272 WT (138 male, 134 female), 318 Het (161 male, 157 female) and 154 KO (70 male, 84 female) mice.

### *Arh2*-KO and -Het MEFs showed increased cell proliferation, colony formation as well as tumorigenesis in nude mice

We showed that *Arh2* mutations induced tumors in mice. To assess the effects of *Arh2* genotype in proliferation, three individual mouse embryonic fibroblasts (MEFs) of each genotype (e.g., WT1-3, Het1-3, KO1-3) were generated from WT and *Arh2*-KO littermates (Supplemental Material and Supplemental Figure 1). Cell proliferation of three individual MEFs among genotypes was monitored for 96-h (Figure 6A). Cell proliferation of all three lines of *Arh2*-KO and -Het MEFs was significantly faster than that of WT MEFs during the 96-h incubation (Figure 6A). To confirm the result of cell proliferation, colony formation assay of MEFs among genotypes was examined (Figure 6B-D). Numbers and size of colonies in all three lines of *Arh2*-KO and -Het MEFs were significantly greater than those of WT MEFs (Figure 6B-D). Thus, *Arh2* mutations increased cell proliferation, and colony formation. To examine the effect of *Arh2* genotype in tumor formation, MEFs among genotypes were injected into athymic nude mice and monitored for 62 days. Three individual MEFs in each genotype were injected subcutaneously into nude mice. Tumor mass in nude mice was quantified three time per week and followed for 62 days. All three lines of *Arh2*-KO and -Het MEFs formed tumor, but not WT MEFs (Figure 6E). These results suggested that *Arh2*-KO and -Het MEFs increased tumorigenesis in nude mice.

**Figure 6.**
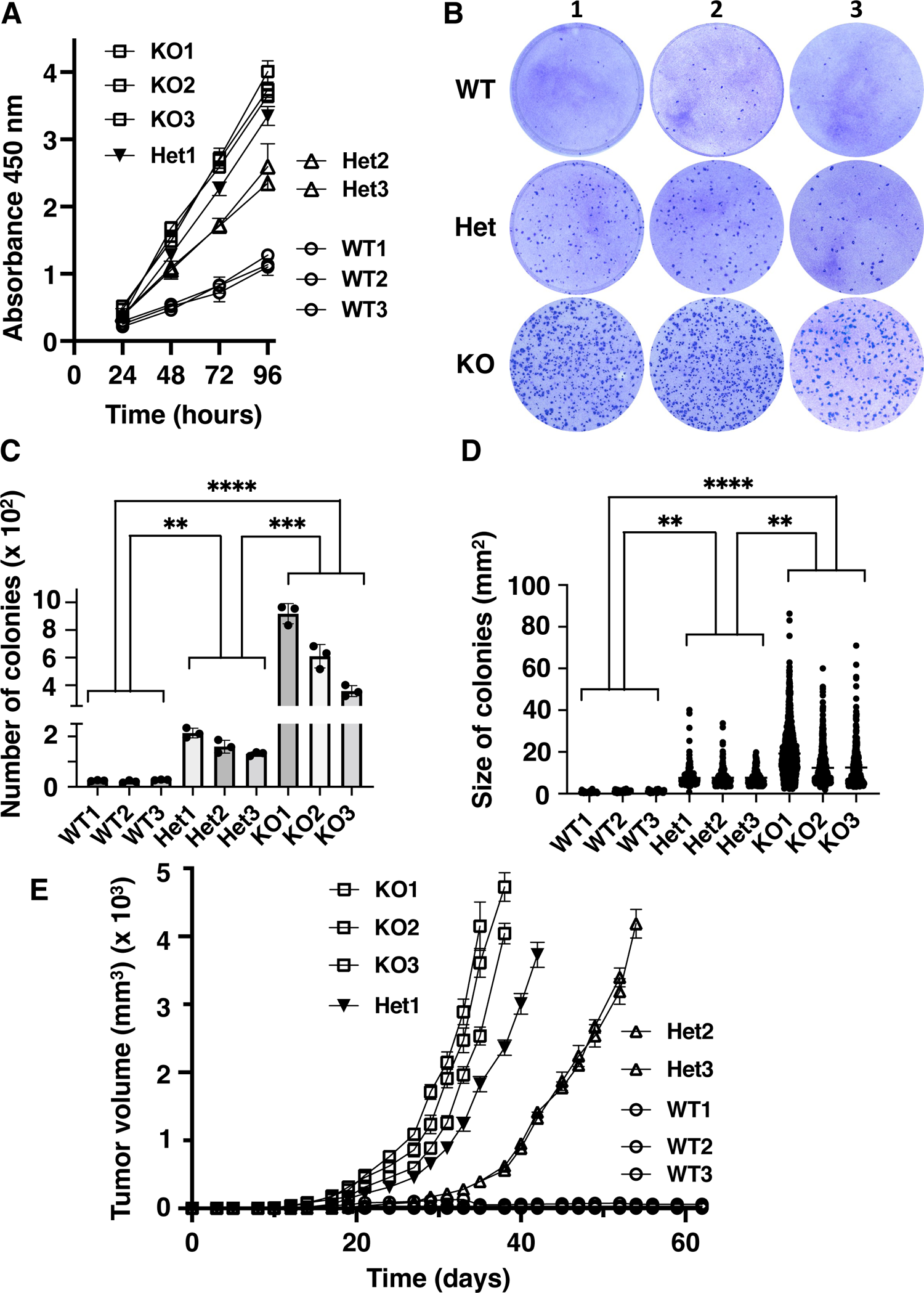
*Arh2*-KO and -Het MEFs showed increased cell proliferation, colony formation, and tumorigenesis in nude mice. (**A**) Effects of *Arh2* genotype in cell proliferation of MEFs. Cell proliferation assay of three individual MEFs in each genotype, wild-type (WT, ○), *Arh2*-heterozygous (Het, ▽, Δ), *Arh2*-knockout (KO, □) were assessed during the 96-days. Data are shown mean ± SEM of values from 4 assays, 4 experiments. Significant difference, \*\*\*\**P* < 0.0001 by two-way ANOVA, Tukey’s multiple comparison tests. Not significant between KO2 vs. KO3, Het2 vs. Het3, WT1 vs. WT2 or WT3, and WT2 vs WT3. (**B**) Representative digital images (100 mm petri-dish) in three individual WT, *Arh2*-Het, and *Arh2*-KO MEFs. Colonies were stained with crystal violet after incubation for 16 days (5% CO_2_ at 37 °C). (**C**) Number of colonies in three individual WT, *Arh2*-Het, and *Arh2*-KO MEFs after incubation for 16 days (5% CO_2_ at 37 °C). Data are shown mean ± SEM of values from duplicates, 3 experiments in each MEFs. Significant difference, \*\**P* < 0.01, \*\*\**P* < 0.001, \*\*\*\**P* < 0.0001 by one-way ANOVA, Tukey’s multiple comparison tests. Not significant between KO2 vs. KO3, among Het1-3 MEFs, and among WT1-3 MEFs. (**D**) Size of colonies (mm^2^) in three individual WT, *Arh2*-Het, and *Arh2*-KO MEFs after incubation for 16 days (5% CO_2_ at 37 °C). Data are shown mean ± SEM of values from duplicates, 3 experiments in each MEFs. Significant difference, \*\**P* < 0.01, \*\*\**P* < 0.001, \*\*\*\**P* < 0.0001 by one-way ANOVA, Tukey’s multiple comparison tests. Not significant between KO2 vs. KO3, among Het1-3 MEFs, and among WT1-3 MEFs. (**E**) Tumor formation in nude mice injected with three individual WT (○), *Arh2*-Het (▽, Δ), *Arh2-*KO (□) MEFs via subcutaneously. Width and length in tumor masses were measured 3 times per week thereafter. Data are shown mean ± SEM of values from 5 nude mice per each MEFs, and two experiments. Tumor growth in nude mice injected with all *Arh2-*KO or all *Arh2*-Het MEFs showed significantly faster than those in nude mice injected with all WT MEFs at \*\*\*\**P* < 0.0001 by two-way ANOVA, Tukey’s multiple comparison tests.

### Mutation of p.L116R, c.347 T>G, exon 2 in *Arh2* gene plays a critical role in tumorigenesis, corresponding to human *ARH2* mutation in stomach adenocarcinoma

We determined *Arh2*-homozygous mutations cause tumorigenesis in mice, and nude mice using MEFs. In addition, *Arh2*-heterozygous with mutation showed results similar to those seen in *Arh2*-homozygous mutation. To determine which functional allele is involved in tumorigenesis, sequence analysis of *Arh2*-Het MEFs, tumor tissues in nude mice injected with *Arh2*-Het MEFs, and spontaneous tumor tissues in *Arh2*-Het mice was investigated. Three individual *Arh2*-Het MEFs (Het1-3), 15 tumor tissues from nude mice injected with Het1-3 MEFs each injected into 5 nude mice, 24 spontaneous tumor tissues (e.g., 5 hepatocellular carcinomas, 5 histiocytic sarcomas, 4 metastatic histiocytic sarcomas in liver, 6 lymphomas, 4 metastatic lymphomas in liver) from *Arh2*-Het mice were subjected to DNA sequence analysis. The list of *Arh2* mutations (e.g., synonymous, substitution-missense) in *Arh2*-Het MEFs, and tumor tissues in nude mice injected with *Arh2*-Het MEFs is shown in Supplemental Table 1, and in spontaneous tumors in *Arh2*-Het mice is shown Supplemental Table 2. Four mutations in exons 1 and 2 of *Arh2* gene were identified from Het1 MEFs and tumor tissues in nude mice injected with Het1-3 MEFs (Supplemental Table 1). Fourteen mutations in exons 1, 2, 3, 4 and 7 of *Arh2* gene were identified from spontaneous tumor tissues in *Arh2*-Het mice (Supplemental Table 2). In contrast, mutations were not observed in Het2 or Het3 MEFs or all WT (WT1-3) MEFs as well as in adjacent non-tumor tissue, surrounding normal tissue, and peripheral blood from the *Arh2-*Het mice. Of interest, 15 of 18 mutations in *Arh2* gene derived from the MEFs and tumor tissues (e.g., Het1 MEFs, tumors in nude mice injected with *Arh2*-Het1 MEFs, spontaneous tumors) corresponded to human *ARH2* genes in human cancers obtained from Catalogue Of Somatic Mutations In Cancer (COSMIC) (https://cancer.sanger.ac.uk/cosmic/gene/analysis?ln=ADPRHL1) (Supplemental Table 3, Supplemental Figure 4 and 5). In the results of the tumor growth curve of nude mice injected with Het1-3 MEFs, those of all (5 of 5) nude mice injected with Het1 MEFs, 2 of 5 nude mice injected with Het2 MEFs, and 2 of 5 nude mice injected with Het3 MEFs showed significantly greater proliferation than those in other 6 nude mice injected with Het2 or Het3 MEFs (Supplemental Figure 4A). Consistently, their tumor volume doubling time was significantly shorter than the other 6 nude mice injected with Het2 or Het3 MEFs (Supplemental Figure 4B). Two mutations of *Arh2* gene (p.A7A, c.21 A>G, exon 1; p.L116R, c.347 T>G, exon 2) in nude mice injected with Het1 MEFs were observed. Of note, those aggressive tumor tissues exhibited the same *Arh2* mutation (p.L116R, c.347 T>G, exon 2) (Supplemental Table 1 and Supplemental Figure 5). Thus, mutations of p.A7A, c.21 A>G, exon 1 and p.L116R, c.347 T>G, exon 2 in *Arh2* gene are associated with aggressive tumor growth in nude mice. Surprisingly, mutation of p.L116R, c.347 T>G, exon 2 in *Arh2* gene corresponded to human *ARH2* mutation in stomach adenocarcinoma obtained from COSMIC, suggesting that mutation of p.L116R, c.347 T>G, exon 2 in *Arh2* gene plays a critical role in tumorigenesis in mice and human (Supplemental Table 3 and Supplemental Figure 5).

### Both genders of *Arh2*-KO and -Het mice displayed increased unexpected deaths and decreased survival rate caused by tumor and inflammation diseases

We showed that *Arh2* mutant mice displayed pathological phenotypes, cardiac dysfunction, and tumorigenesis. To determine the effect of *Arh2* mutations on survival rate, unexpected morbidity, and mortality, *Arh2* mice were monitored for 24-month. Mice who experienced unexpected deaths or met endpoint criteria of animal protocol were subjected to pathological evaluation. The survival rate of both genders of *Arh2*-KO and -Het mice was significantly decreased during the 24-month observation period (Figure 7A). The list of unexpected deaths in both genders of mice among *Arh2* genotypes is shown in Supplemental Table 4. Incidence in mice of unexpected death classified with tumor or non-tumor in both genders of *Arh2*-KO (male, 100%; female, 94%) and -Het mice (male, 88%; female, 58.6%) was significantly greater than those of WT (male, 23.2%; female, 23.1%) mice (Figure 7B, Supplemental Table 4). Of note, about half of unexpected deaths was classified as non-tumor in *Arh2*-mutant mice (Figure 6B). Incidence of unexpected deaths of non-tumor in *Arh2*-KO (male, 50%; female, 57.1%) and -Het (male, 41%; female, 30.6%) mice were significantly greater than those of WT (male, 15.9%; female, 17.2%) mice (Figure 7C). To examine what types of non-tumor origin affected unexpected deaths in *Arh2*-mutant mice, unexpected non-tumor deaths were classified as inflammation, non-inflammation, and congenital diseases (within 3-month-old) in *Arh2* mice (Figure 7D-F).

**Figure 7.**
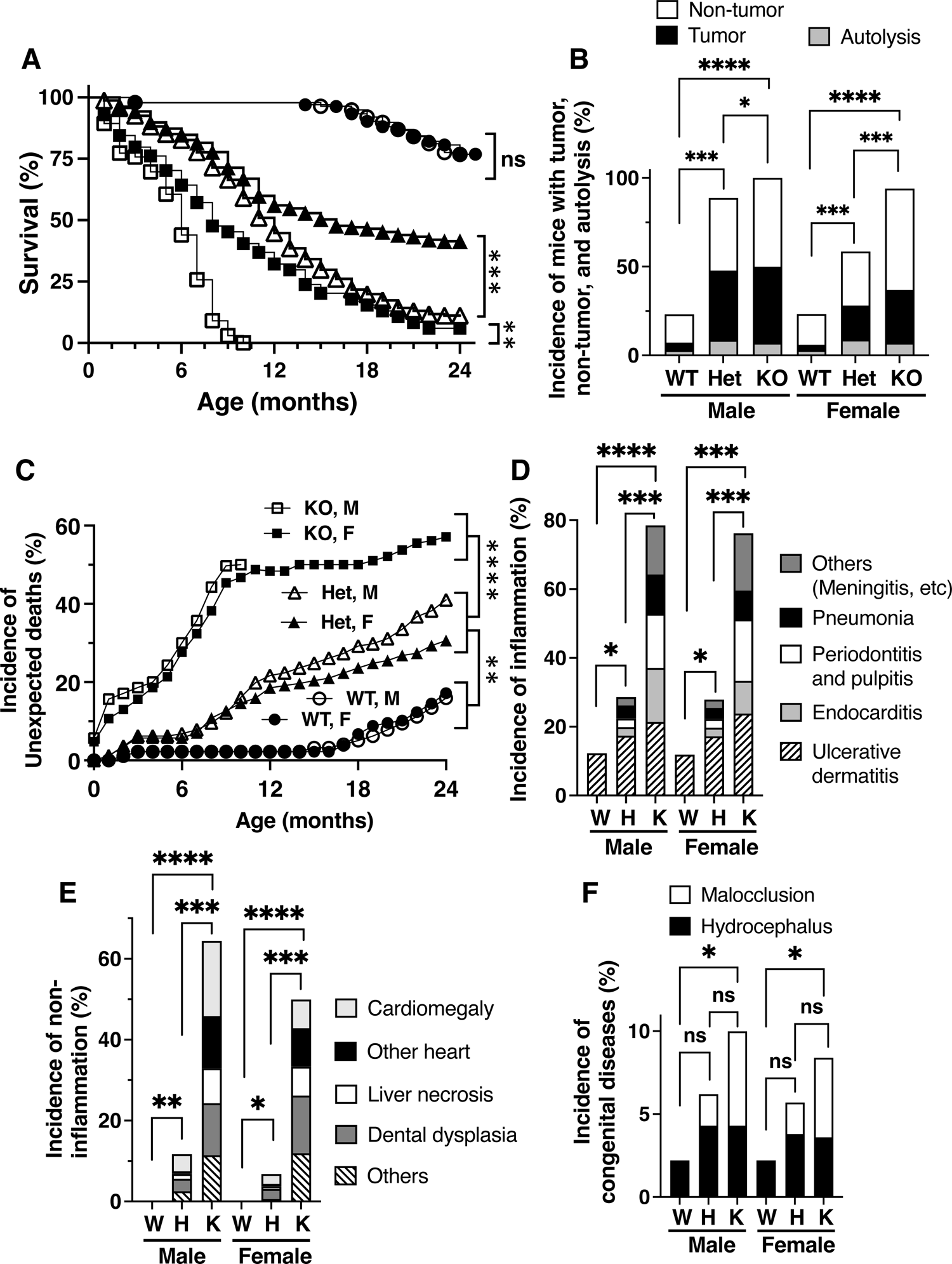
*Arh2*-KO and -Het mice showed increased unexpected deaths and decreased survival rate. (**A**) Survival rate in male and female mice among *Arh2* genotypes during the 24-month observation. Survival of 272 WT (138 male, 134 female), 318 *Arh2*-Het (161 male, 157 female) and 154 *Arh2*-KO (70 male, 84 female) mice were monitored during the 24-month. Significant difference, *Arh2*-KO male vs WT male; \*\*\*\**P* < 0.0001, *Arh2*-Het male vs WT male \*\*\*\**P* < 0.0001; *Arh2*-KO male vs *Arh2*-Het male, \*\**P* < 0.01; *Arh2*-KO female vs *Arh2*-Het female, \*\**P* < 0.001. Survival analysis in each pair was performed by Logrank Mantel-Cox and Gehan-Breslow-Wilcoxon tests. (**B**) Incidence of unexpected deaths in male and female mice among *Arh2* genotypes during the 24-month observation. Tumor (black), non-tumor (white) and autolysis (gray) are classified. Significant difference, \**P* < 0.05, \*\*\**P* < 0.001, \*\*\*\**P* < 0.0001 by one-way ANOVA, Tukey’s multiple comparison tests. (**C**) Comparison the cumulative number of unexpected deaths derived from non-tumors by *Arh2* genotypes and age during the 24-month observation. The % of unexpected death in WT mice (male ○, female •), *Arh2*-Het mice (male Δ, female Δ), *Arh2*-KO mice (male □, female ▀) were indicated. Significant difference, *Arh2*-KO male mice vs WT male mice, \*\*\*\**P* < 0.0001; *Arh2*-KO female mice vs WT female mice, \*\*\*\**P* < 0.0001; *Arh2*-Het male mice vs WT male mice, \*\*\*\**P* < 0.0001; *Arh2*-Het female mice vs WT female mice, \*\*\*\**P* < 0.0001; *Arh2*-KO male mice vs Het male mice, \*\*\*\**P* < 0.0001; *Arh2*-KO female mice vs Het female mice, \*\*\*\**P* < 0.0001 by one-way ANOVA, Tukey’s multiple comparison tests. Not significant between *Arh2*-KO male mice vs. *Arh2*-KO female mice, *Arh2*-Het male mice vs. *Arh2*-Het female mice, WT male mice vs. WT female mice. (**D**) Incidence of inflammation derived from non-tumors of the unexpected deaths in *Arh2* mice among genotypes. Non-tumors derived from unexpected deaths were subjected to histopathological evaluation. Ulcerative dermatitis (borders), endocarditis (light gray), periodontitis (white), pneumonia (black), others (meningitis), glomerulonephritis, otitis, and metritis (dark gray) were observed in *Arh2*-KO and -Het mice. Only ulcerative dermatitis was observed in WT mice. Significant difference, \**P* < 0.05, \*\*\**P* < 0.001, \*\*\*\**P* < 0.0001 by one-way ANOVA, Tukey’s multiple comparison tests. (**E**) Incidence of non-inflammation derived from non-tumors of the unexpected deaths in *Arh2* mice among genotypes. Non-tumors derived from unexpected deaths were subjected to histopathological evaluation. Cardiomegaly (light gray), other heart diseases including cardiac infarction, thrombus and metaplasia in aortic valve, and cardiac artery mineralization (black), liver necrosis (white), dental dysplasia (dark gray) and others including spleen telangiectasia, bilateral end stage crystal in kidney, and stomach ulcer (borders) were observed in *Arh2*-KO and -Het mice. Only ulcerative dermatitis was observed in WT mice. Significant difference, \**P* < 0.05, \*\*\**P* < 0.001, \*\*\*\**P* < 0.0001 by one-way ANOVA, Tukey’s multiple comparison tests. Any non-inflammation was not observed in WT mice. Significant difference, \**P* < 0.05, \*\**P* < 0.01, \*\*\**P* < 0.001, \*\*\*\**P* < 0.0001 by one-way ANOVA, Tukey’s multiple comparison tests. (**F**) Incidence of congenital diseases derived from non-tumors of the unexpected deaths in *Arh2* mice among genotypes. Non-tumors derived from unexpected deaths were subjected to histopathological evaluation. Hydrocephalus (black) and malocclusion (white) were observed in *Arh2*-KO and -Het mice. Only hydrocephalus was observed in WT mice. Significant difference, \**P* < 0.05, by one-way ANOVA, Tukey’s multiple comparison tests.

The variety of inflammation (e.g., endocarditis, pneumonia, periodontitis, pulpitis, ulcerative dermatitis), non-inflammation (e.g., cardiomegaly, liver necrosis, dental dysplasia), and congenital disease (e.g., hydrocephalus, malocclusion) were observed in *Arh2*-KO and -Het mice (Figure 6D-F, Supplemental Figure 6 and Supplemental Table 5). Incidence of inflammation in both genders of *Arh2*-KO (male, 78.4%; female, 76.2%) and -Het (male, 28.6%; female, 27.9%) mice was significantly greater than those in WT mice (male, 12.3%; female, 11.9%) (Figure 6D and Supplemental Figure 6A-D). Incidence of non-inflammation in both genders of *Arh2*-KO (male, 64.5%; female, 50.1%) and -Het (male, 11.7%; female, 6.8%) mice was significantly greater than those in WT mice (male, 0%; female, 0%) (Figure 6E and Supplemental Figure 6E-G). Consistent with the results seen in histopathology (Figure 1), incidence of cardiomegaly in both genders of *Arh2*-KO mice has significantly increased compared to those of WT mice (Supplemental Figure 5E). Incidence of congenital diseases in both genders of *Arh2*-KO mice (10% of male, 8% of female) was significantly increased relative to those of WT (2% male, 2% female) mice (Figure 6F). These results suggested that *Arh2*-KO and -Het mice showed increased unexpected deaths and decreased survival rate caused by tumor and non-tumor diseases.

### *Arh2* homozygous mutation caused partial embryonic lethality

We showed that ARH2 plays an important role in survival, preventing the development of multiple abnormalities. To determine the effect of ARH2 in pups and embryos development, genotype rate of pups after birth and of embryo among genotypes was assessed. Expected genotype percentages of pups or embryos are 25% of WT, 50% of *Arh2*-Het, and 25% of *Arh2*-KO (Supplemental Figure 7A). However, genotype percentages of *Arh2*-KO pups (20%: 70 males, 84 females) and *Arh2*-Het pups (43%: 161 males, 157 females) were decreased (Supplemental Figure 7A). Instead of decreased % of *Arh2*-KO and -Het pups, genotype percentages of WT pups (37%: 138 males, 134 females) were increased (Supplemental Figure 7A). Consistent with the results seen in genotype percentages of pups after birth, genotype percentages in both 13-14 days (E13-14) and 17-18 days (E17-18) embryos of *Arh2*-KO (E13-14:19%, E17-18:19%) were decreased more than expected (Supplemental Figure 7B). Genotype percentages of WT embryos (E13-14: 36%, E17-18:35%) and of *Arh2*-Het embryos (E13-14: 45%, E17-18: 46%) were increased (Supplemental Figure 7B). Thus, *Arh2* deletion resulted in partial embryonic lethality in inbred heterozygous intercrosses.

### Analysis of ADP-ribose binding site on predicted human ARH2 compared with crystal structure of human ARH1 and ARH3

In contrast to ARH1 and ARH3, ARH2 does not exhibit any enzymatic activities (19, 25). To understand the basis of the difference in ARH-family enzymes, we analyzed ADP-ribose binding site by comparing predicted structure of human ARH2 compared with crystal structures of human ARH1 and ARH3. Structure comparison of AlphaFold predicted human ARH2/ADPRHL1 (white, UniProt #Q8NDY3) and human ARH1 (brown, PDB ID: 6g28) with bound ADP-ribose and Mg^2+^ were shown (Supplemental Figure 8A and B). Predicted human ARH2 structure (white) is similar to the ARH1 structure (Supplemental Figure 8A). However, Phe130 (Cys in ARH1, Asn in ARH3) and Glu304 (Asp in ARH1/3) substitution in ARH2 seem interfere with ADP-ribose and metal binding, respectively (Supplemental Figure 8B). Arg98 (Ala in ARH1) in ARH2 seems to block entrance of ADP-ribosylated protein substrates (e.g., ADP-ribose-Arg). In addition, the sequence alignment shows that ARH2 does not have Mg-coordinating and substrate-binding residues that are present in ARH1 and ARH3 in the sequence alignment (Supplemental Figure 9). Thus, our sequence and structural analysis support the lack of enzymatic activities in ARH2.

## Discussion

*Arh2*-KO mice exhibited cardiac dysfunction with cardiomegaly and muscle dysfunction assessed by histological analysis and physiological testing (e.g., MRI, dobutamine stress test by echocardiography, motor function tests) (Figure 1-3). *Arh2*-Het mice showed results similar to those seen in *Arh2*-KO mice except for cardiomegaly (Figure 1-3). *Arh2*-KO and -Het mice developed spontaneous tumors consistent with the result seen in mutant MEFs that caused tumor formation in nude mice (Figure 4, 5). DNA sequence analysis of *Arh2*-Het MEFs and heterozygous tumors derived from nude mice injected with *Arh2*-Het MEFs revealed that 13 of 18 mutations corresponded to human *ARH2* mutations seen in cancers in COSMIC database (Supplemental Figure 5 and Supplemental Table 1-3). Of interest, the L116R mutation in *Arh2* gene was associated with aggressive tumor growth in nude mice. The identical L116R mutation was found in the COSMIC database in a human stomach adenocarcinoma (Supplemental Figure 5 and Supplemental Table 1 and 3). Both genders of *Arh2*-KO and -Het mice showed increased unexpected deaths and decreased survival rate caused by tumor and non-tumor diseases during a 24-month observation (Figure 6). Thus, *Arh2* is an essential gene for survival by maintaining cardiac function, and preventing tumor formation and non-tumor diseases.

### *Arh2* homozygous and heterozygous mutations developed cardiac dysfunction and motor dysfunction

Cardiomegaly with cardiac fibrosis was observed in both genders of *Arh2*-KO mice in an age-dependent manner (Figure 1). Consistently, incidence of cardiomegaly during the 24-month observation in *Arh2*-KO mice has significantly increased compared to the WT mice (Supplemental Figure 5). *Arh2*-KO mice showed decreased cardiac contractility (Figure 2). In addition, *Arh2*-Het mice showed decreased ejection fraction at high-dose dobutamine stress assessed by echocardiography (Figure 3). Similar results to cardiac effects, both genders of *Arh2*-KO and -Het male mice showed skeletal muscle dysfunction (Figure 4). To support our data, Stuart J. Smith *et al*. reported that ARH2 in *Xenopus* embryos is associated with heart chamber outgrowth/function and acts on muscle actin filament assembly in the forming ventricle (17, 28). Since ARH2 is mainly expressed in heart and skeletal muscle (15–17, 28), our findings showed that ARH2 plays an essential role in heart and muscle function.

### *Arh2* homozygous and heterozygous mutations developed tumorigenesis

*Arh2*-KO and -Het mice exhibited developed spontaneous tumors during the 24-month observation (Figure 5). Although prostate tumor and uveal melanoma, which were previously reported in families with *ARH2* mutation (29–31), were not observed in spontaneous tumors of *Arh2*-KO and -Het mice, *Arh2*-KO and -Het mice developed different types of tumors (e.g., adenocarcinoma, hepatocellular carcinoma, lymphoma, histiocytic sarcoma) (Figure 4). Uncontrolled cell proliferation is a hallmark of cancer cells (32). Indeed, *Arh2*-KO and -Het MEFs showed increased cell proliferation and colony formation (Figure 6A-D). Consistently, *Arh2*-KO and -Het MEFs showed increased tumorigenesis in nude mice (Figure 6E). Thus, *Arh2* homozygous and heterozygous mutations participate in development of tumorigenesis in mice.

### Mutation of p.L116R, c.347 T>G, exon 2 in *Arh2* gene plays a pivotal role in tumorigenesis in mice and human

We identified 18 mutations of *Arh2* gene in tumorigenesis derived from *Arh2*-Het MEFs and heterozygous tumor tissues (e.g., tumor formed in nude mice after injecting *Arh2*-Het MEFs, spontaneous tumor in *Arh2*-Het mice) (Supplemental Table 1 and 2). Of note, 13 of 18 mutations corresponded to human *ARH2* mutations in various cancers in COSMIC database. Study of tumor growth and tumor volume doubling time in nude mice after injecting Het1-3 MEFs revealed that mutations of p.A7A, c.21 A>G, exon 1 and p.L116R, c.347 T>G, exon 2 in *Arh2* gene were associated with aggressive tumor growth, which derived from quickly growing tumors in nude mice after injecting Het1 MEFs (Supplemental Figure 4 and Supplemental Table 1). In addition, other quickly growing tumors derived from Het2 or Het3 MEFs in nude mice exhibited the same mutations of p.L116R, c.347 T>G, exon 2 in *Arh2* gene (Supplemental Table 1). Surprisingly, mutation of p.L116R, c.347 T>G, exon 2 in *Arh2* gene corresponded to human *ARH2* mutation in stomach adenocarcinoma in COSMIC database (Supplemental Table 1 and 3). Thus, mutation of p.L116R, c.347 T>G, exon 2 in *Arh2* gene plays critical role in tumorigenesis in mice and human.

### *Arh2* homozygous and heterozygous mutations increased unexpected deaths and decreased survival rate caused by tumorigenesis and non-tumor diseases

During the 24-month observation, *Arh2*-KO and -Het mice displayed increased unexpected deaths and decreased survival rate (Figure 7). Regarding survival rate, both genders of *Arh2*-KO mice died within 10 months (Figure 7). In addition, *Arh2* deletion resulted in partial embryonic lethality in inbred heterozygous intercrosses, suggesting that ARH2 plays an essential role in survival and embryo development (Figure 7 and Supplemental Figure 7). Of note, about half of unexpected deaths was caused by tumors in *Arh2* mutant mice (Figure 5B), thus tumorigenesis is the main cause of unexpected deaths in *Arh2* mutant mice. In addition, we showed that *Arh2*-KO and -Het mice exhibited variety of abnormalities, including (e.g., endocarditis, pneumonia, periodontitis, pulpitis, ulcerative dermatitis), non-inflammation diseases (e.g., cardiomegaly, liver necrosis, dental dysplasia), and congenital disease (e.g., hydrocephalus, malocclusion), leading to unexpected deaths (Figure 7). Indeed, endocarditis, pneumonia, cardiomegaly, liver necrosis, and hydrocephalus are related to cause of death (33–37). Periodontitis, pulpitis, and those comorbidities increased the risk of endocarditis due to infection or cardiac dysfunction (e.g., myocardial infarction, stroke, peripheral vascular disease) (38–40). Dental dysplasia and malocclusion might lead to malnourishment and eventual premature death as feeding becomes more difficult (41–43). Consistent with the results seen in cardiomegaly of *Arh2*-KO mice (Figure 1), incidence of cardiomegaly classified in non-inflammation in *Arh2*-KO mice was significantly increased relative to those of WT mice (Supplemental Figure S6E). In addition, 31.5% of causes of unexpected deaths were non-inflammatory heart (Figure 7D and Supplemental Table 5), suggesting that heart non-inflammation is the second cause of unexpected death. Thus, homozygous and heterozygous of *Arh2* mutations increased not only tumorigenesis, but also multiple non-tumor diseases (e.g., inflammation, non-inflammation, congenital diseases) caused unexpected deaths and decreased survival rate.

### ARH2 shares similarities and identities of ARH1

We showed that ARH2 does not have the Mg^2+^-binding amino acids of the catalytic sites analyzed with structural-based amino acid sequences and predicted human ARH2 structure compared with human ARH1 and ARH3 (Supplemental Figure 8 and 9), but its structure was found to be similar to ARH1. Of note, human and mouse ARH1 amino acid was 45-47% identity and 66-68 similarity of human and mouse ARH2 (25), which can explain structural similarity with both predicted ARH2 and crystalized ARH1 structures (Supplemental Figure 8). Similar to ARH2 in phenotypes, *Arh1*-KO mice developed tumorigenesis and cardiac dysfunction with cardiomyopathy (44, 45). To investigate mechanism or signaling pathway how ARH2 is related to cardiac dysfunction tumorigenesis, Omics (e.g., RNA sequence, proteomics) analysis are further needed.

In summary, our mouse model with *Arh2* homozygous and heterozygous mutations have presented the crucial role of ARH2 in heart/muscle function, tumorigenesis, and inflammatory/non-inflammatory/congenital diseases. The knowledge of the role of ARH2 provides a better understanding of the role of this protein in pathogeneses of human diseases.

## Materials and Methods

### Generation of *Arh2* KO mice and expression of ARH2 in *Arh2* mice

We generated *Arh2* knockout (KO) mice using CRISPR/Cas9 gene editing targeting the first exon of the *Arh2* gene (Supplemental Figure 1A), resulting in the deletion of 338 base pairs containing part of the 5’ UTR and part of the coding region. As shown in Supplemental Figure S1B, the PCR products of the wild-type (WT) allele and mutant allele are 660 bp and 322 bp, respectively. The absence of ARH2 protein was confirmed by Western blotting analysis (Supplemental Figure 1C and D). No immunoreactive protein was detected in heart and skeletal muscle of *Arh2* KO mice using anti-ARH2 antibody (see Materials and Methods). Amounts of 39-kDa ARH2 expression in *Arh2* heterozygotes were significantly less in lysates of heart or skeletal muscle than those in *Arh2* WT mice mouse (Supplemental Figure S1C, S1D). In wild-type mice as well as humans, Northern blotting analyses (Supplemental Figure S2) showed that the *Arh2* gene is primarily expressed the heart and skeletal muscle. A relatively lower level of expression was detected in mouse whole embryos, which was likely contributed by the cardiac and muscle tissues within the embryo.

### Animal Study

All mouse studies were performed in accordance with the NIH Guide for the Care and Use of Laboratory animals and with the approval of the NHLBI Animal Care and Use Committee under protocols H-0127R5 and H-0172R5. *Arh2*-knockout (KO) mice were backcrossed for 10 generations with mice having a C57BL6J (Jackson Laboratory) background. Animal facility, which housed in *Arh2* mice, was free of mouse parvovirus, but *Helicobacter* species and mouse norovirus were present. After pups were weaned from breeders (heterozygous male and female), wild-type (WT), heterozygous (HT) and KO mice were maintained in the same cage, but mice were separated based on gender. Mouse genotypes were confirmed by PCR.

### Generation of *Arh2* knockout mouse line

The *Arh2* knockout mouse line was generated using CRISPR/Cas9 (46) (Wang et al, 2013). Briefly, two sgRNAs (Arh2-1:TGCAGATGTTTCCATAGCCG; Arh2-2: CGACAACACCATCATGCACA) were designed to cut shortly downstream of the translation initiation codon (ATG) of the mouse *Arh2* gene. These sgRNAs were made using ThermoFisher’s custom in vitro transcription service. The sgRNAs (20ng/ul each) were co-microinjected with Cas9 mRNA (50 ng/ul, purchased from Trilink Biotechnologies) into the cytoplasm of zygotes collected from B6D2F/J F1 hybrid mice (JAX Stock No. 100006). Injected embryos were cultured overnight in M16 medium (MilliporeSigma) in a 37 °C incubator with 6% CO_2_. The next morning, embryos that reached 2-cell stage of development were implanted into the oviducts of pseudopregnant surrogate mothers. Offspring born to the foster mothers was genotyped by PCR amplification of the targeted region using primer pairs (forward: 5’-gactcctttgctggagagcca-3’; and reverse: 5’-ggctgctgtcccctaaagaca-3’). The PCR bands were then sequenced using Sanger sequencing for confirming desired deletions. All procedures for generating the *Arh2* CRISPR/Cas9 mice were performed in the NHLBI Transgenic Core Facility.

### Collection of *Arh2*-KO pups after birth and embryos

After backcrossed with C57BL6J (B6J) mice and *Arh2* heterozygous mice, *Arh2* mice were interbred to generate litters with *Arh2* wild-type (WT), heterozygous (Het), and knockout (KO) pups and embryos from heterozygous intercrosses. Genotypes of pups and embryos were confirmed by PCR.

### Magnetic resonance imaging (MRI)

Cardiac Magnetic Rresonance Imaging (MRI) was performed in a 7-Tesla MRI system with ParaVision software (Bruker, Billerica, MA). Mice were anesthetized with 2-3% isoflurane and placed in an MRI coil, while monitoring ECG, respiratory rate, and body temperature (SA Instruments, Stony Brook, NY). Magnevist (Bayer Healthcare, Whippany, NJ) diluted 1:10 with sterile saline solution, was administered IP at 0.1 to 0.6 mmol/kg of mouse body weight, 5-10 min before MRI. Following initial images for localization, CINE MRI images were acquired sequentially using a cardiac- and breath-gated, flow-compensated gradient echo method (GEFC). The sequence parameters were TE (echo time) = 3.4ms with a 10ms repetition time between frames, FOV (field of view) = 30 x 30 mm for short axis images or 48 x 28 mm, for long axis images, slice thickness = 1 mm, and matrix = 256 x 256. Image data consisted of 8-10 short axis slices that covered the heart from base to apex, and two perpendicular long-axis slices. MRI was performed in the Mouse Imaging Facility (MIF).

### Baseline and dobutamine stress test by echocardiography

Baseline and dobutamine stress test by echocardiography in *Arh2* mice was performed using the Vevo 2100 imaging system (FUJIFILM VisualSonics, Bothell, WA), with a MS-400 transducer (FUJIFILM VisualSonics), with a center operating frequency of 30 MHz. Mice were anesthetized with isoflurane (2%) via nose cone during echocardiography and dobutamine infusions and placed on a heated platform with a rectal temperature probe and electrocardiography (ECG) leads. After baseline echocardiography, low- and high-dose dobutamine (10, 40 µg/kg/min) solution (0.625 mg/ml dobutamine in saline containing 5% dextrose) were administered by intravenous infusion via a tail vein catheter using syringe pump (Cole Parmer) at constant rate infusion. When heart rate was stabilized at approximately 500 beats/min after low-dose dobutamine (10 µg/kg/min) infusion, then, high-dose (40 µg/kg/min) was administered. 2D images were obtained for multiple views of the heart, and M-mode images of the left ventricle were collected from the parasternal short axis view at the level of the midpapillary muscles. From the M-mode images, LV systolic and diastolic posterior and anterior wall thicknesses, as well as end-systolic and end-diastolic internal LV chamber dimensions (LVIDs, LVIDd) were measured using the leading-edge method. The percentage of ejection fraction (%EF) and percentage of fractional shortening (%FS) were calculated from measurements by the machine’s internal software.

### Mouse exercise stress test (Treadmill test)

At 7-8 months of age, *Arh2* mice were tested for exercise capacity using a Columbus Instruments rodent treadmill (Model Eco-6M), set at a 10-degree incline. Testing protocol was as follows: 10 min with belt speed at 10 m/min, 12 m/min for five min, 15 m/minute for three minutes. Then, the belt speed was increased incrementally by 1.8 m/min every three min until the mouse became exhausted. Total time, distance, maximum speed, and work were recorded at the time of exhaustion, defined as when mice were unable to continue running without repeatedly falling back onto the shock grid at the back of the treadmill.

### In Vivo Assay of muscle contraction strength (Isometric torque test)

In vivo muscle contractility was measured with the plantar flexor muscles (gastrocnemius and soleus) of lower limb, using an Aurora Scientific Inc model 1300A. Mice were placed supine on a thermostatically controlled table under inhaled anesthesia, 4-5% isoflurane for induction, using induction chamber for less than a min, and then, up to 2% isoflurane via a nosecone for maintenance, both at 1 to 1.5 L/min 100% O2. The knee was kept stationary, and the foot was firmly fixed to a footplate, which was connected to the shaft of the motor. The foot was aligned at 90° to the tibia. Muscle contraction (ankle plantar flexion) was elicited by placing subcutaneous electrodes to directly stimulate the plantar flexor muscles (gastrocnemius and soleus). A force frequency protocol was performed by applying a series of stimulation frequencies from 25 to 250 Hz at 25-Hz increments with a pause of 1 min between stimuli, and maximal isometric torque (mN-m) was measured at full tetanic contraction at each frequency. To account for differences in body size among 7–8-month-old *Arh2* mice, maximal isometric torque was normalized to body weight (kg), which is shown as specific torque (mN-m/kg).

### Accelerating-rotarod test

This test assesses motor coordination, balance, and equilibrium in mice. Each mouse was tested on the Rotamex 5 rotarod apparatus (Columbus Instruments, Columbus, OH) and first acclimated for 2 days before the test, at a static rod (60sec, 3trials, 10 min rest between each) and steady speed of 4 rpm, (maximum 3 min/trial, 3 trials/training, with 10 min resting time between each trial). For the test, the rotarod was set at an accelerating mode (4–40 rpm, 5 min), and time (seconds) for the mouse to fall from the rotarod was recorded. For each mouse, the test was repeated 3 times with 1 h rests between each trial.

### Anti-ARH2 Antibodies

Rabbits were immunized with a peptide (CKVTFPDNYDAEERDKT) representing amino acids 243-258 (exons 5-6) of mouse ARH2 with cysteine added at the N-terminus to facilitate coupling to keyhole limpet hemocyanin. Antibodies were purified by Yenzyme Inc (Brisbane, CA) from sera of two rabbits, using a peptide affinity column.

### Cell culture

Mouse embryonic fibroblasts (MEFs) were generated from 14.5-days embryo from breeding set of *Arh2* heterozygous mice. *Arh2* genotypes of all MEFs were confirmed by PCR. MEFs were cultured in Dulbecco’s modified Eagle’s medium (DMEM) containing 10% fetal bovine serum (FBS, Life Technologies), 100 U/ml penicillin, and 100 μg/ml streptomycin (Life Technologies) in a humidified atmosphere containing 5% CO_2_ at 37 °C.

### Cell proliferation assay

*Arh2* MEFs, 3 x 10^3^ /well, were seeded on 96-well plates, incubated for 24, 48, 72, or 96 h in a humidified atmosphere containing 5% CO_2_ at 37 °C. Cell number was assessed using a Cell Counting Kit-8 (Dojindo) by measuring absorbance at 450 nm, according to the manufacturer’s instructions. Absorbance was measured using SpectraMax M5 Microplate Reader (Molecular Devices).

### Colony formation assay

*Arh2* MEFs, 3 x 10^3^ were seeded on 100 mm petri dishes and incubated for 16 days in a humidified atmosphere containing 5% CO_2_ at 37 °C for colony formation. After removing media and washing each dish with 10 mL 0.9% saline, colonies on each dish were fixed with 10 mL 10% neutral, buffered-formalin solution for 30 minutes, and then stained 10 mL 0.01% (w/v) crystal violet (Sigma) in distilled water for 30-60 min. After washing off excess crystal violet with distilled water and drying the dishes, digital images of the dishes with colonies were obtained using a stereomicroscope. Colony number and size on each dish were quantified using ImageJ (version 1.53k).

### Tumorigenicity in nude mice

Six- to eight-week-old athymic female (JAX, homozygous Foxn1nu/Foxn1nu, Stock #007850) each 5 mice were injected subcutaneously to three individual WT, *Arh2*-Het, and *Arh2*-KO MEFs (passage 6 of 1 x 10^6^ cells in 0.15 mL DMEM per site). Tumor volume (mm^3^) was measured 3 times per week using electronic digital calipers from Fowler & NSK) and calculated as length x width^2^ x 0.5 (47). Mice were euthanized with CO_2_ when tumors reached a length of 2.0 cm or growth of tumor was observed into the skin or on 62 days after injection. Tumors were subjected to histopathologic examination.

### SDA-PAGE and Western blotting

ARH2 protein was detected by Western blotting. Protein (20 μg) from extracted cell lysate was subjected to SDS-PAGE (4-12% NuPAGE gel, Life Technologies), transferred to nitrocellulose membranes. The membrane was blocked with 5% non-fat dry milk (Bio-Rad) in TBS-T (Tris-Buffered-Saline with 0.05% tween-20) at 4 °C overnight or room temperature for 1 h, and then incubated with primary antibody at 4 °C, overnight, or room temperature for 1 h. The membrane was washed with TBS-T and then incubated with a secondary antibody of IRDye 800CW goat anti-rabbit (LI-COR) in blocking buffer at room temperature for 1 h. The membrane protein was detected by an Odyssey Imaging Systems (LI-COR).

### Histopathology

Histopathological tissues including tumors from *Arh2* mice were evaluated using tissue cross-sections stained with hematoxylin and eosin (HE). Myocardial fibrosis was evaluated using tissue cross-sections stained with HE or Masson’s trichrome stain for collagen-rich fibrotic regions. Percentage of fibrotic area was calculated by MetaMorpha software using Masson’s trichrome-stained sections. Histology sections were evaluated by NIH DVR rodent pathological section or NHLBI Pathological Core. Histological slides were subsequently digitalized using a digital slide scanner (NanoZoomer, Hamamatsu Photonics KK, Hamamatsu, Japan). Digital images were taken by NDP.view2 software (Hamamatsu Photonics KK, Hamamatsu, Japan).

### COSMIC database search for *ARH2* gene

Data in Supplementary Table S3 are based on the COSMIC database https://cancer.sanger.ac.uk/cosmic (COSMIC v97 release).

### Statistics

ANOVA (One-way or Two-way) analysis of variance (repeated-measures analysis of variance methods using Bonferroni’s and Tukey’s multiple-comparison tests) were used. All data were analyzed for statistical significance using GraphPad Prism 9 software. All statistical tests were considered significant at the 0.05 level.

## Supporting information

ARH2 MRI Movie

## Acknowledgments

We thank Rodney L. Levine for valuable advice. We thank the staff at Diagnostic and Research Service Branch of Division of Veterinary Resources for necropsies and preparation of slides. I.K.K. was supported by National Institutes of Health (R01 GM141226 to I.K.K.).

## Author contributions

J.K. observed *Arh2* mice, designed, acquired, analyzed, and interpreted data, and drafted and wrote the manuscript; S.Y. designed, analyzed, and interpreted data, and drafted and wrote the manuscript; H.I.E. helped designed experiments and analyzed data; S.O. confirmed enzymatic activity and performed Northern blot analysis; Z.X.Y. and V.H. assisted with preparation of histopathological specimens, and pathological evaluation; C.L. and J.K. generated *Arh2*-knockout mice. D.A.S., A.N., and M.P. performed echocardiography, treadmill, rotarod, and torque tests. M.J.L. performed MRI analysis. I.K.K and M.M. analyzed ARHs structural comparisons. J.M conceptualized, designed and interpreted data, drafted, and reviewed the manuscript. All the authors have reviewed and approved the manuscript.

## Supplemental Figure legends

**Supplemental Figure 1.**
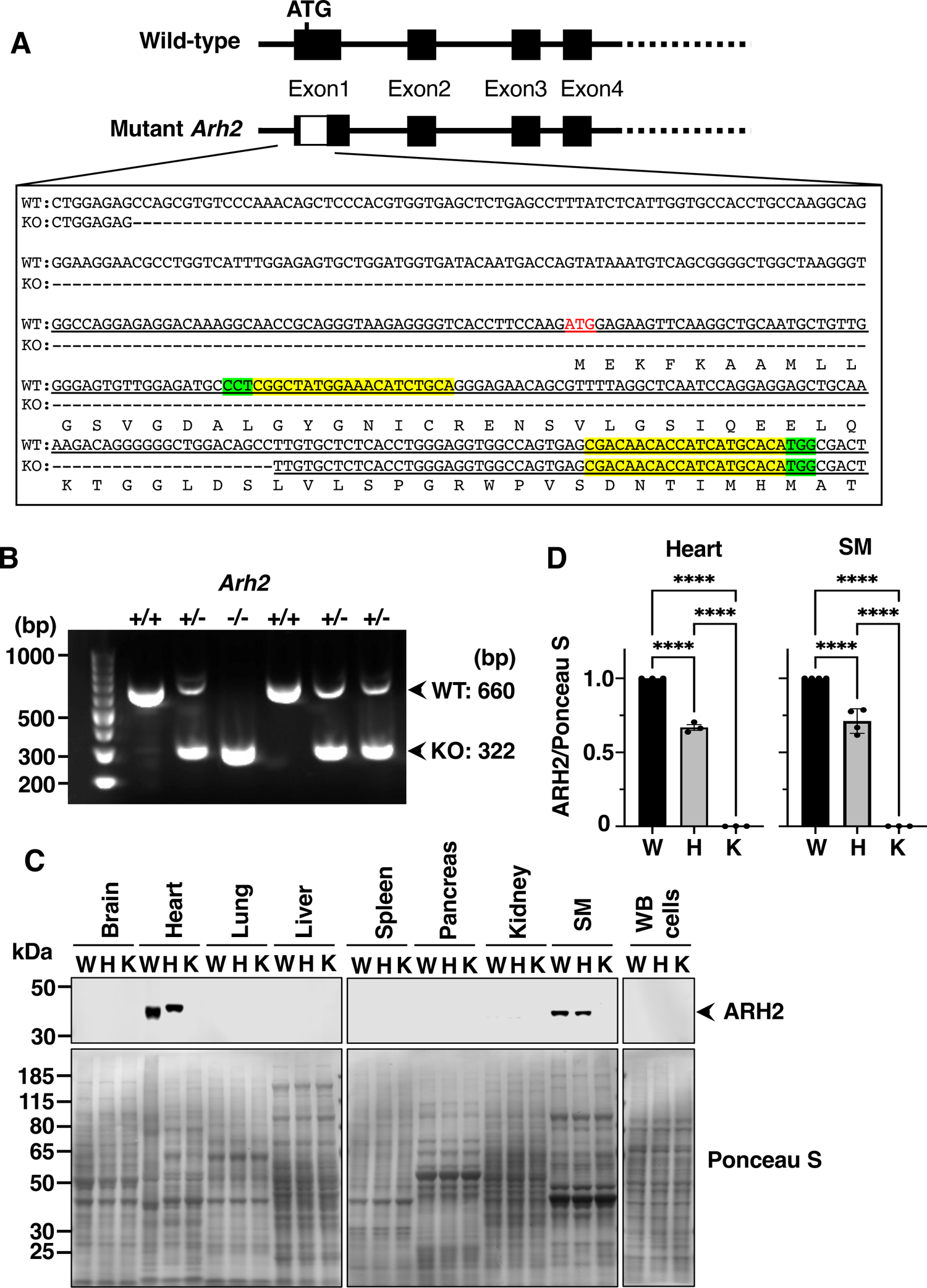
Generation of *Arh2*-KO and -Het mice. (**A**) The targeted site of *Arh2* locus for deletion. Diagram of CRISPR-mediated 338bp deletion in the *Arh2* locus (dashes, open box) with exons 1 to 4 shown as boxes while the promoter and introns are represented by a line. The deletion includes a part of the 5’ UTR (untranslated region) and the beginning part of the coding sequence (including the ATG start codon, red). CRISPR sgRNA binding sites are highlighted in yellow and their PAMs (protospacer adjacent motif) are highlighted in green. (**B**) Representative image of mouse genotyping by PCR amplification of mouse tail genomic DNA. *Arh2* wild-type (+/+) and homozygous/knockout (-/-) mice clearly yield the 660bp WT band and 322bp mutant band, respectively, while heterozygous (+/-) mice yield both bands. (**C**) Representative images of Western blotting analysis of ARH2 protein expression in tissues. Wild-type (W), *Arh2*-heterozygous (H), and *Arh2*-knockout (K) mice are indicated. Skeletal muscle is abbreviated as SM and white blood cells are abbreviated as WB. Upper blots show immunoblot using anti-ARH2 peptide antibody (ARH2 murine amino acid 243-258, exon 5-6, CKVTFPDNYDAEERDKT). Lower blots show Ponceau S stained. Arrow indicates ARH2 (39 kDa). (**D**) Densitometric analysis of ARH2 expression normalized with total protein in heart and skeletal muscle (SM) of *Arh2* wild-type (W) and heterozygous (H) and knockout (K) mice. Data was presented as mean ± S.E.M. in duplicate and three experiments. Significant difference, **** *P* < 0.0001 by one-way ANOVA, Tukey’s multiple comparison tests. Data are generated using Odyssey (LI-COR) and ImageJ software.

**Supplemental Figure 2.**
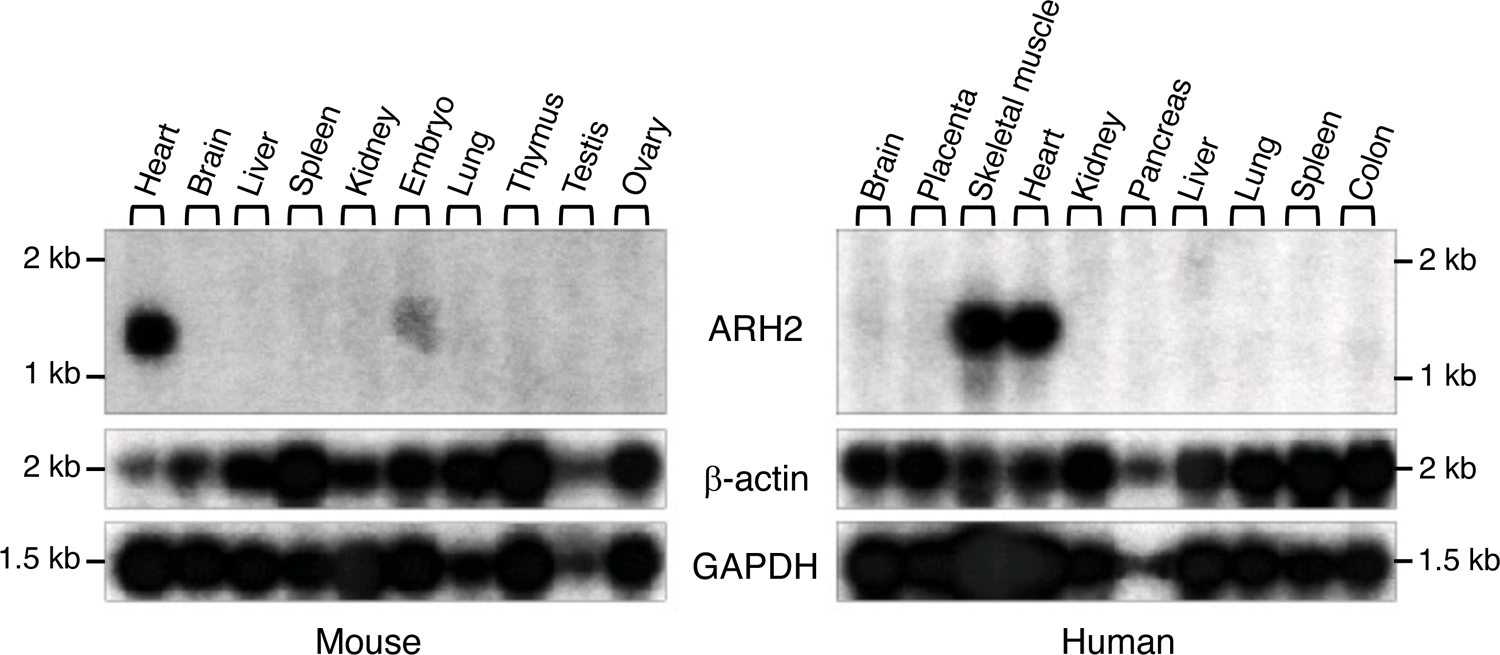
*ARH2* mRNA in mouse and human tissues. A hybridization of mouse poly(A) RNA with ARH2 cDNA probe. Northern blot of poly(A) RNA (2μg) from mouse tissues (Thermofisher) was hybridized at 42 °C in 30 ml of hybridization buffer (Thermofisher) overnight with 25 ng of ^32^P-labeled ARH2 cDNA. In a separate experiment, ^32^P-labeled (10 ng) glyceraldehyde-3-phosphate dehydrogenase and β-actin cDNAs were hybridized to the blot as a loading control. Positions of RNA standards are shown on the left. Data shown reflect hybridization of mouse poly(A) RNA with ARH2 cDNA probe, representative of results of Northern analyses of human and mouse tissues performed four times.

**Supplemental Figure 3.**
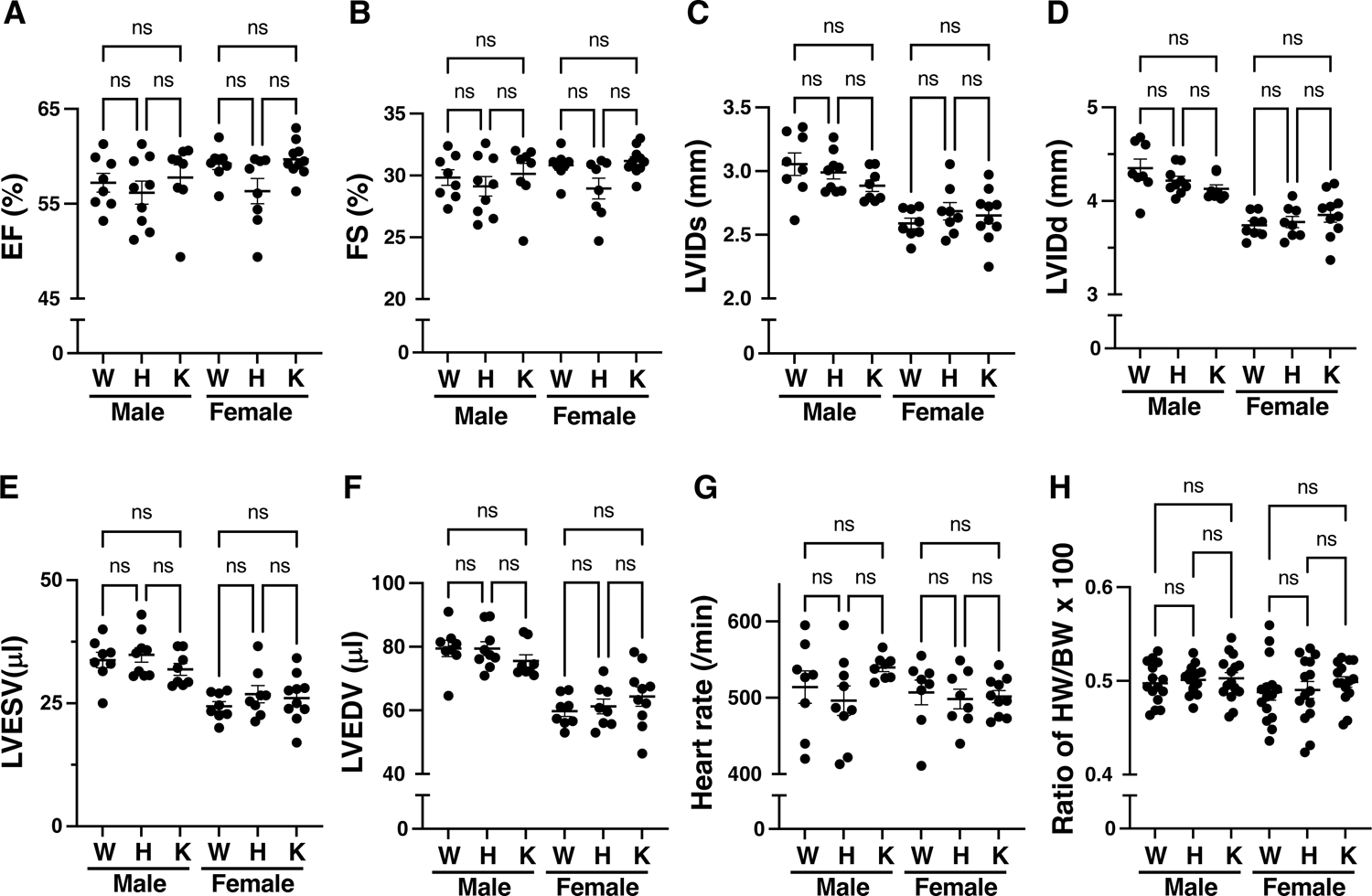
3-4-month-old *Arh2* mice among genotypes showed normal cardiac contractility assessed by echocardiography and ratio of heart weight to body weight. (**A-G**) Echocardiography analysis at baseline in the 3-4-month-old of male and female mice among *Arh2* genotypes. Wild-type (W), *Arh2*-heterozygous (H), and *Arh2*-knockout (K) mice are indicated. Cardiac echocardiography parameters: ejection fraction (%EF, **A**), fractional shortening (%FS, **B**), left ventricular internal diameter (mm) at systole (LVIDs, **C**), and at diastole (LVIDd, **D**), and left ventricular chamber volume (μL) at systole (LVESV, **E**) and at diastole (LVEDV, **F**), and heart rate (/min; **G**). *n* = 8-10. Data were compared by one-way ANOVA Tukey’s multiple comparison test. There is no significant difference among genotypes regardless of gender. ns: no significant. (**H**) Ratio of heart weight (HW) to body weight (BW) in the 3-4-month-old male and female mice among genotypes. *n* = 15. Data were compared by one-way ANOVA, Tukey’s multiple comparison tests. There is no significant difference among genotypes regardless of gender. ns: no significant.

**Supplemental Figure 4.**
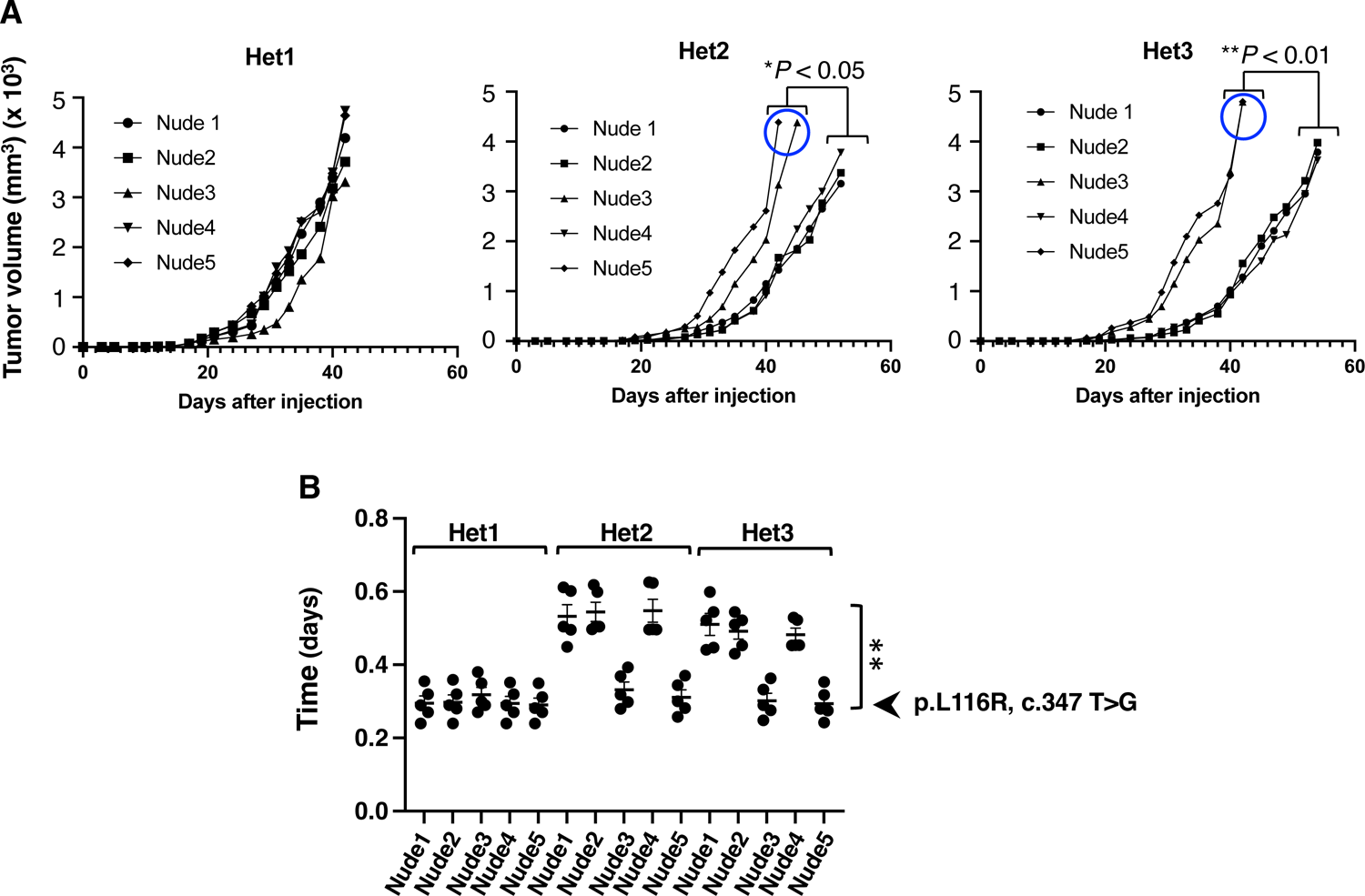
Effects of *Arh2* gene mutations on tumor growth in nude mice. (**A**) Tumor growth curve in nude mice injected with three individual *Arh2*-Het MEFs (Het1, Het2, and Het3) via subcutaneously during the 62 days observation. Those of all (5 of 5) nude mice injected with Het1 MEFs, 2 of 5 nude mice injected with Het2 MEFs (blue circle), and 2 of 5 nude mice injected with Het3 MEFs (blue circle) showed significantly greater proliferation than those in other 6 nude mice injected with Het2 or Het3 MEFs. Data are shown mean ± SEM of values from 5 nude mice per each MEFs, and two experiments. Significant difference, \**P* < 0.05, \*\**P* < 0.01 by one-way ANOVA, Tukey’s multiple comparison tests. (**B**) Tumor volume doubling time (TvDT) in tumors injected with three individual *Arh2*-Het MEFs (Het1, Het2, and Het3). TvDT of all (5 of 5) nude mice injected with Het1 MEFs, 2 of 5 nude mice injected with Het2 MEFs (blue circle), and 2 of 5 nude mice injected with Het3 MEFs (blue circle) showed significantly greater than those in other 6 nude mice injected with Het2 or Het3 MEFs. Significant difference, \*\**P* < 0.01 by one-way ANOVA, Tukey’s multiple comparison tests. Those aggressive tumor tissues exhibited the same *Arh2* mutation (p.L116R, c.347 T>G, exon 2) (Supplemental Table 1 and Supplemental Figure 5).

**Supplemental Figure 5.**
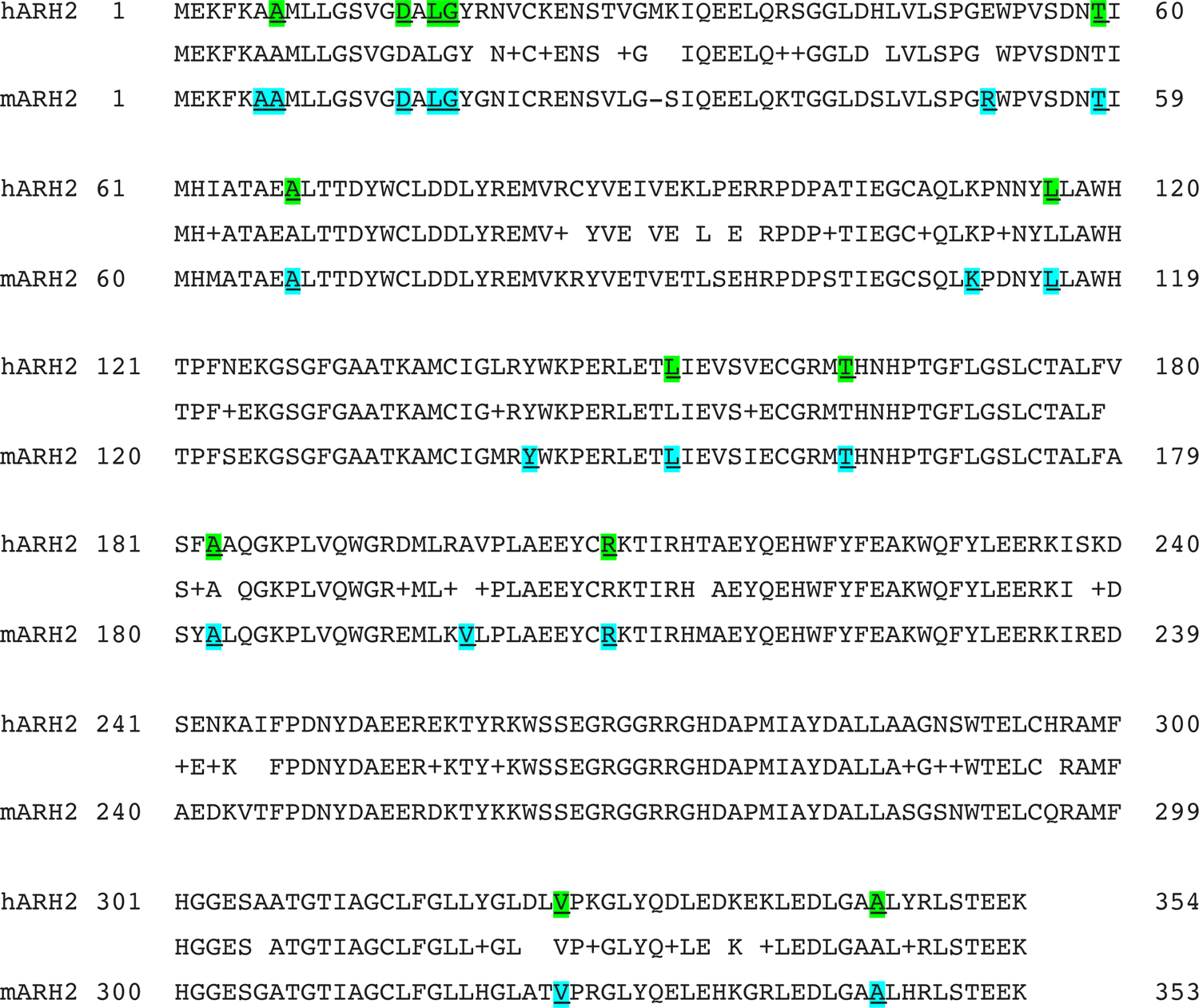
ARH2 mutation site in human and murine cancer with amino acids pairwise alignments. Eighty five percent identities and 93% similarities (middle line) were seen between human ARH2 (hARH2, upper line) and murine ARH2 (mARH2, lower line) in ARH2 protein alignments. Highlight shows ARH2 mutation sites in human (green) and murine (cyan blue) cancer.

**Supplemental Figure 6.**
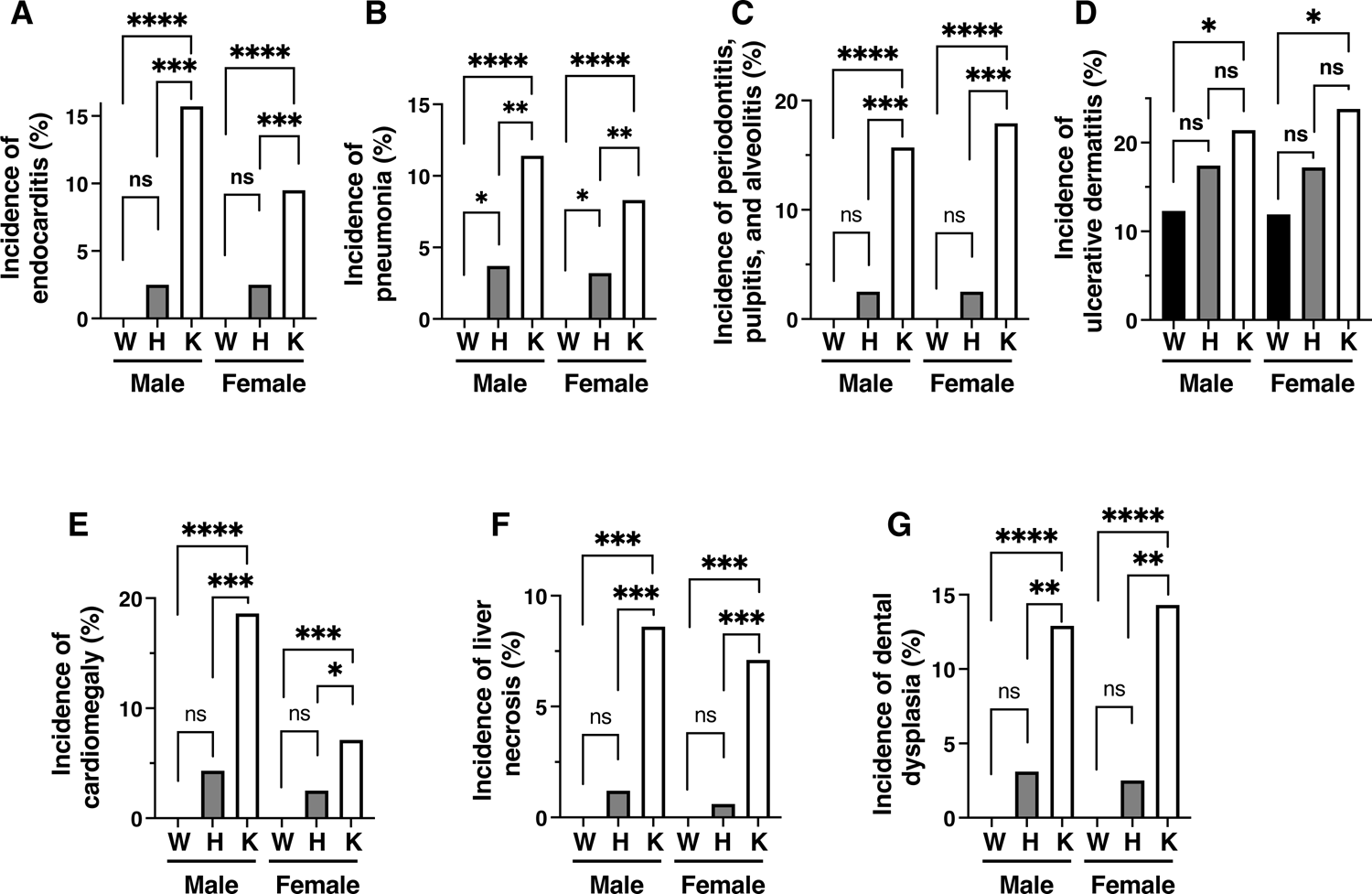
*Arh2* KO mice showed increased inflammation and non-inflammation comorbidities. (**A**-**D**) Incidence of unexpected deaths with inflammation in male and female among *Arh2* genotypes during the 24-month observation. Wild-type (W), *Arh2*-heterozygous (H), and *Arh2*-knockout (K) mice are indicated. Histopathological evaluation showed endocarditis (**A**), pneumonia (**B**), periodontitis including pulpitis and alveolitis (**C**), ulcerative dermatitis (**D**) in *Arh2* mice. Significant difference, \**P* < 0.05, \*\**P* < 0.01, \*\*\**P* < 0.001, \*\*\*\**P* < 0.0001 by one-way ANOVA, Tukey’s multiple comparison tests. (**E**-**G**) Incidence of unexpected deaths with non-inflammation in male and female among *Arh2* genotypes during the 24-month observation. Histopathological evaluation showed cardiomegaly (**E**), liver necrosis (**F**), dental dysplasia (**G**) in *Arh2* mice. Significant difference, \**P* < 0.05, \*\**P* < 0.01, \*\*\**P* < 0.001, \*\*\*\**P* < 0.0001 by one-way ANOVA, Tukey’s multiple comparison tests.

**Supplemental Figure 7.**
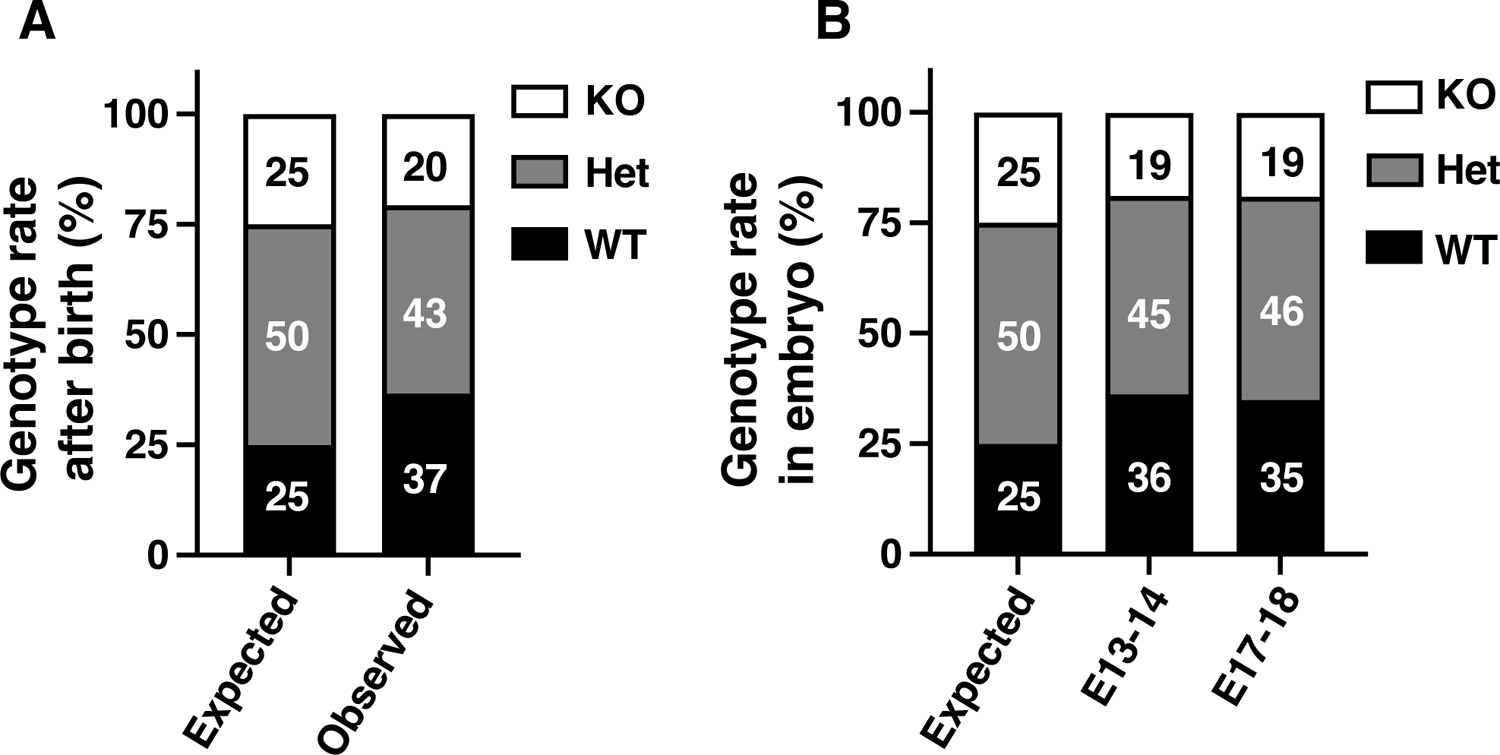
Effects of *Arh2* genotype on number of pups after birth and embryo. (**A**) Genotype rate of pups after birth among genotypes. Bar graph showed percent of genotype ratio in WT (black), *Arh2*-Het (gray), and *Arh2*-KO (white) pups at 3-4 weeks old of age. Bar graph of expected showed a mendelian ratio (1:2:1) of genotypes when heterozygous x heterozygous breeding. Bar graph of observed showed an actual ratio of *Arh2* genotype when *Arh2* heterozygous x *Arh2* heterozygous breeding. Data are shown values from 272 WT (138 male, 134 female), 318 *Arh2*-Het (161 male, 157 female) and 154 *Arh2*-KO (70 male, 84 female) mice. Significant difference, expected *Arh2*-Het vs. observed *Arh2*-Het, \**P* < 0.05; expected *Arh2*-KO vs. observed *Arh2*-KO, \**P* < 0.05; expected WT vs. observed WT, \*\**P* < 0.01 by one-way ANOVA, Tukey’s multiple comparison tests. (**B**) Genotype rate of embryo among genotypes. Bar graph showed percent of genotype ratio in WT (black), *Arh2*-Het (gray), and *Arh2*-KO (white) embryo. Bar graph of expected showed a mendelian ratio (1:2:1) of genotypes when heterozygous x heterozygous breeding. Bar graph of embryo 13-14 days (E13-14) or embryo 17-18 days (E17-18) showed an actual ratio of *Arh2* genotype when *Arh2* heterozygous x *Arh2* heterozygous breeding. Data are shown values of embryos and genotype from 15 pregnant heterozygous female mice. Significant difference, expected WT vs. E13-14 WT or E17-18 WT, \**P* < 0.05; expected *Arh2*-KO vs. E13-14 *Arh2*-KO or E17-18 *Arh2*-KO, \**P* < 0.05 by one-way ANOVA, Tukey’s multiple comparison tests. There is no significant difference between expected *Arh2*-Het and E13-14 *Arh2*-Het or E17-18 *Arh2*-Het.

**Supplemental Figure 8.**
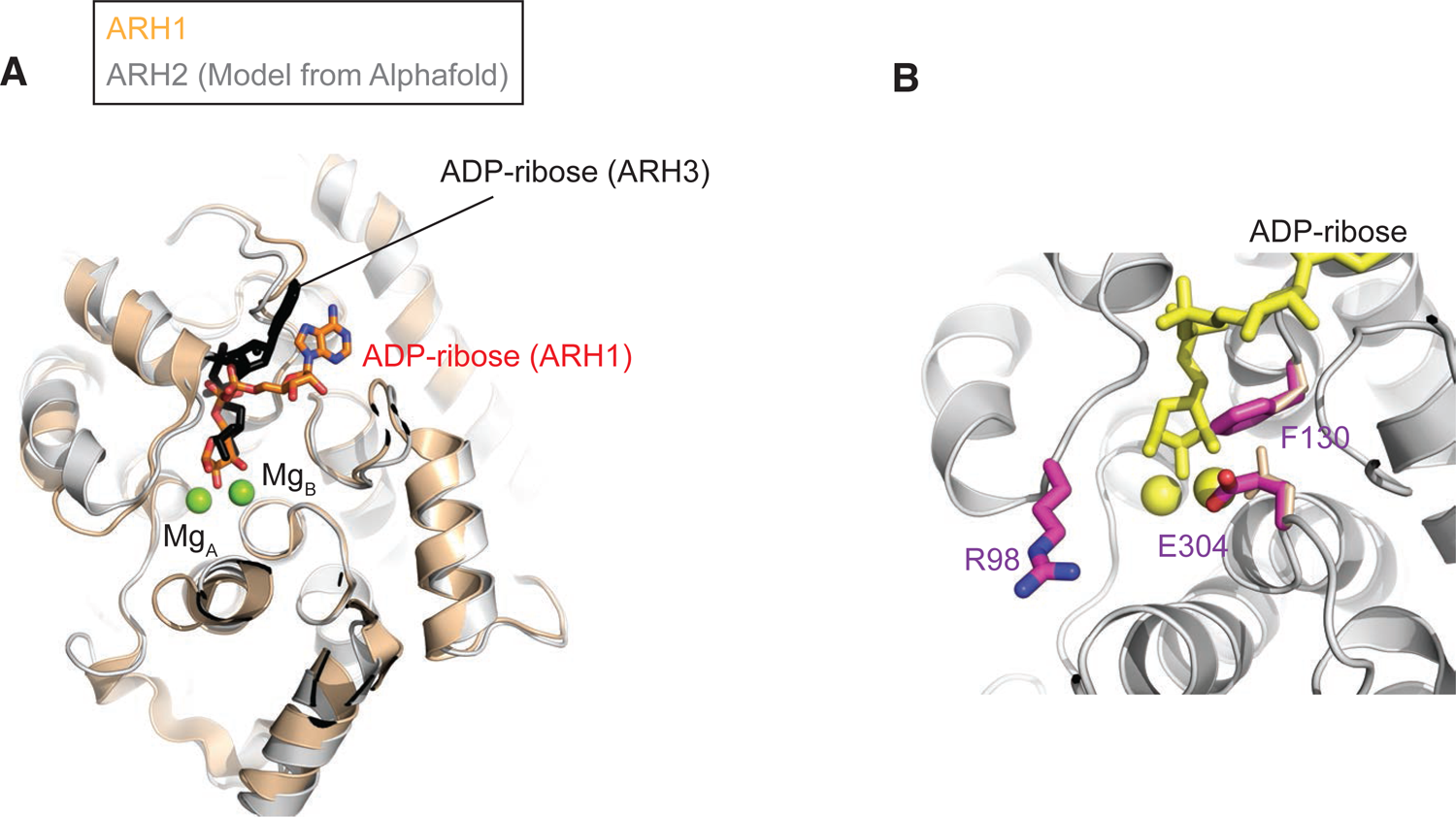
Structural comparison of human ARH2 model with structures of human ARH1 and ARH3 reveals the basis of the lack of enzymatic activity in ARH2. (**A**) Structure comparison of AlphaFold predicted human ARH2/ADPRHL1 (white, UniProt #Q8NDY3) (https://www.uniprot.org/uniprotkb/Q8NDY3/entry) and human ARH1 (brown, PDB ID: 6g28) with bound ADP-ribose and Mg^2+^. (**B**) AlphaFold predicted human ARH2/ADPRHL1 (white, UniProt #Q8NDY3) with ADP-ribose (ARH1). F130 (Cys in ARH1, Asn in Arh3) and E304 (Asp in ARH1/3) substitution seem to interfere with ADP-ribose and metal binding, respectively. R98 (Ala in ARH1) seem to block entrance of ADP-ribosylated protein (e.g., ADP-ribose-Arg).

**Supplemental Figure 9.**
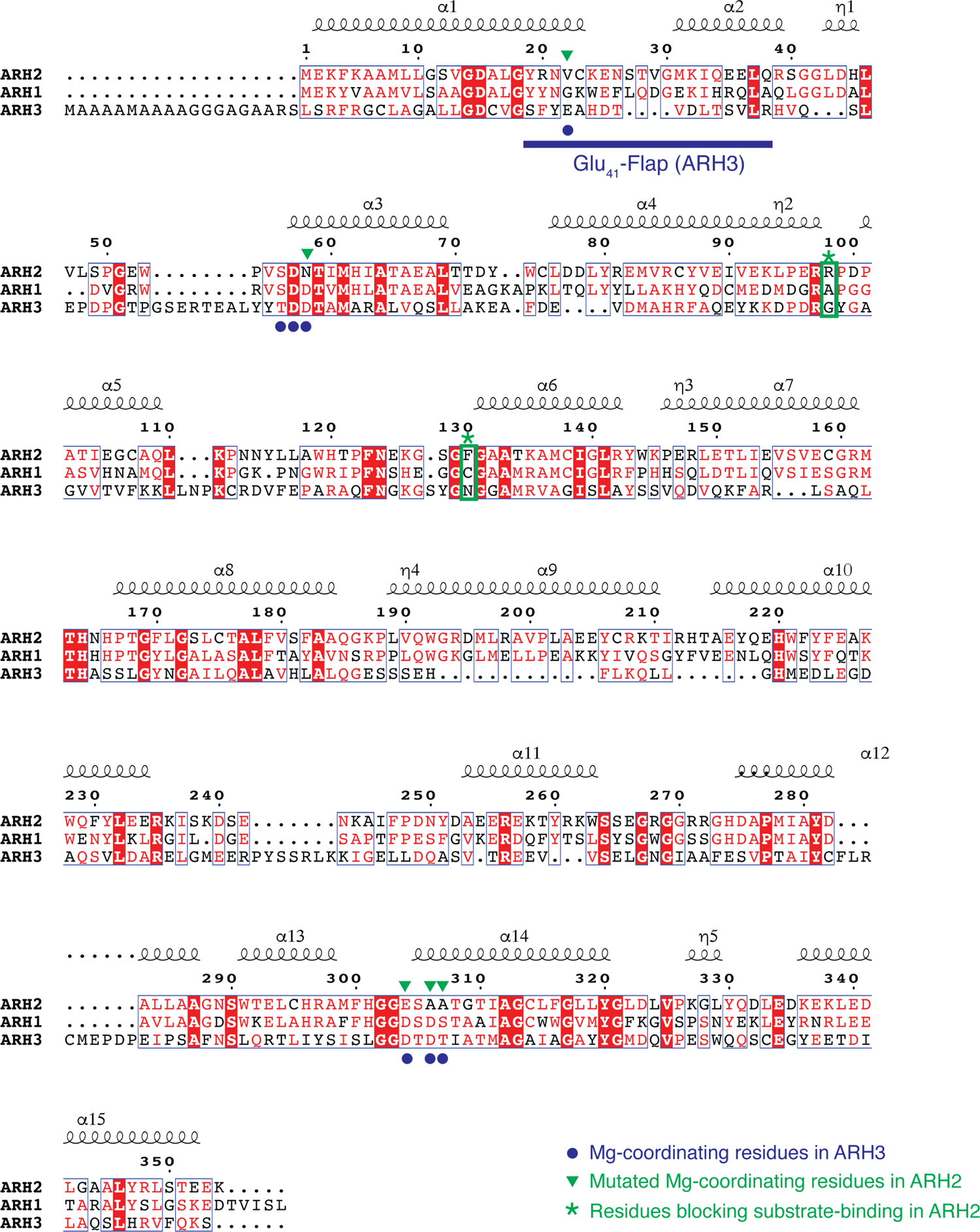
Structure-based alignment of human ARHs. Amino acid and secondary structure elements of human ARH2 compared with human ARH1 and ARH3. Blue dots indicated Mg^2+^ coordinating residues in ARH3. Green arrowheads indicated mutant Mg^2+^-coordinating residues in ARH2. Asterisks on R98 and F130 showed residues blocking substrate-binding in ARH2.

---

**Supplemental Table 1.**
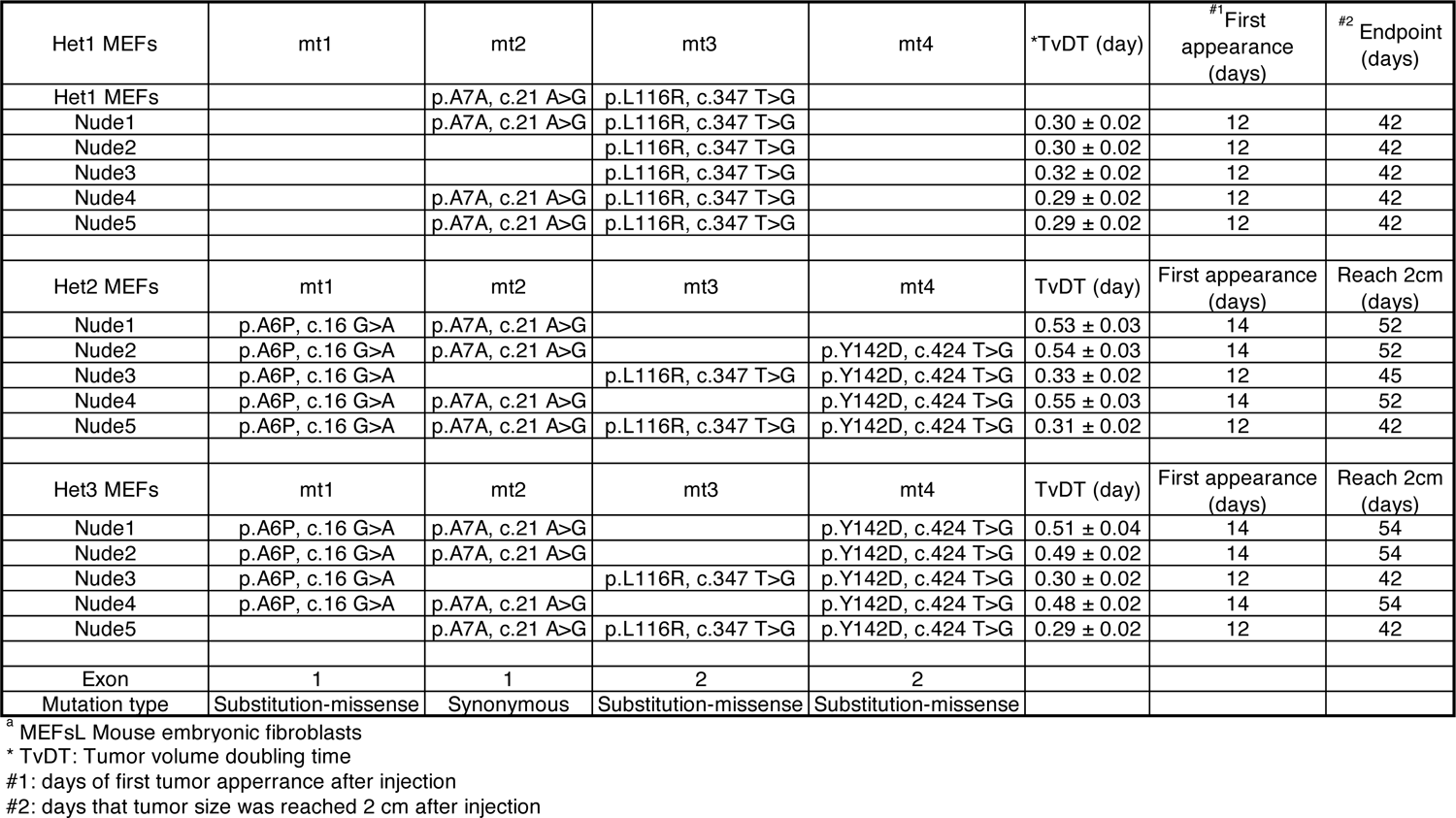
*Arh2* mutations in *Arh2*-Het MEFs and tumors in nude mice injected with *Arh2*-Het MEFs

| Het1 MEFs | mt1 | mt2 | mt3 | mt4 | *TvDT (day) | <sup>#1</sup> First appearance (days) | <sup>#2</sup> Endpoint (days) |
| --- | --- | --- | --- | --- | --- | --- | --- |
| Het1 MEFs |  | p.A7A, c.21 A>G | p.L116R, c.347 T>G |  |  |  |  |
| Nude1 |  | p.A7A, c.21 A>G | p.L116R, c.347 T>G |  | 0.30 ± 0.02 | 12 | 42 |
| Nude2 |  |  | p.L116R, c.347 T>G |  | 0.30 ± 0.02 | 12 | 42 |
| Nude3 |  |  | p.L116R, c.347 T>G |  | 0.32 ± 0.02 | 12 | 42 |
| Nude4 |  | p.A7A, c.21 A>G | p.L116R, c.347 T>G |  | 0.29 ± 0.02 | 12 | 42 |
| Nude5 |  | p.A7A, c.21 A>G | p.L116R, c.347 T>G |  | 0.29 ± 0.02 | 12 | 42 |
| Het2 MEFs | mt1 | mt2 | mt3 | mt4 | TvDT (day) | First appearance (days) | Reach 2cm (days) |
| Nude1 | p.A6P, c.16 G>A | p.A7A, c.21 A>G |  |  | 0.53 ± 0.03 | 14 | 52 |
| Nude2 | p.A6P, c.16 G>A | p.A7A, c.21 A>G |  | p.Y142D, c.424 T>G | 0.54 ± 0.03 | 14 | 52 |
| Nude3 | p.A6P, c.16 G>A |  | p.L116R, c.347 T>G | p.Y142D, c.424 T>G | 0.33 ± 0.02 | 12 | 45 |
| Nude4 | p.A6P, c.16 G>A | p.A7A, c.21 A>G |  | p.Y142D, c.424 T>G | 0.55 ± 0.03 | 14 | 52 |
| Nude5 | p.A6P, c.16 G>A | p.A7A, c.21 A>G | p.L116R, c.347 T>G | p.Y142D, c.424 T>G | 0.31 ± 0.02 | 12 | 42 |
| Het3 MEFs | mt1 | mt2 | mt3 | mt4 | TvDT (day) | First appearance (days) | Reach 2cm (days) |
| Nude1 | p.A6P, c.16 G>A | p.A7A, c.21 A>G |  | p.Y142D, c.424 T>G | 0.51 ± 0.04 | 14 | 54 |
| Nude2 | p.A6P, c.16 G>A | p.A7A, c.21 A>G |  | p.Y142D, c.424 T>G | 0.49 ± 0.02 | 14 | 54 |
| Nude3 | p.A6P, c.16 G>A |  | p.L116R, c.347 T>G | p.Y142D, c.424 T>G | 0.30 ± 0.02 | 12 | 42 |
| Nude4 | p.A6P, c.16 G>A | p.A7A, c.21 A>G |  | p.Y142D, c.424 T>G | 0.48 ± 0.02 | 14 | 54 |
| Nude5 |  | p.A7A, c.21 A>G | p.L116R, c.347 T>G | p.Y142D, c.424 T>G | 0.29 ± 0.02 | 12 | 42 |
| Exon | 1 | 1 | 2 | 2 |  |  |  |
| Mutation type | Substitution-missense | Synonymous | Substitution-missense | Substitution-missense |  |  |  |

<sup>a</sup> MEFsL Mouse embryonic fibroblasts
\* TvDT: Tumor volume doubling time
#1: days of first tumor apperance after injection
#2: days that tumor size was reached 2 cm after injection

**Supplemental Table 2.** *Arh2* mutations in spontaneous tumors of *Arh2*-Het mice.

| Tumors | Mouses *Arh2* mutation | Amino acid mutation | # CDS mutation | Exon | Sample number | Mutation type |
| --- | --- | --- | --- | --- | --- | --- |
| Histiocytic sarcoma 1 | mt5 | p.D15N | c.43 G>A | 1 | 1 | Substitution-missense |
|  | mt11 | p.K110K | c.330 A>G | 2 | 1 | Synonymous |
| Histiocytic sarcoma 2 | mt5 | p.D15N | c.43 G>A | 1 | 1 | Substitution-missense |
| Histiocytic sarcoma 3 | mt12 | p.L151F | c.451 C>T | 3 | 1 | Substitution-missense |
| Histiocytic sarcoma 4 | mt15 | p.V198E | c.593 T>A | 4 | 1 | Substitution-missense |
| Histiocytic sarcoma 5 | mt17 | p.V324I | c.970 G>A | 7 | 1 | Substitution-missense |
| Metastatic histiocytic sarcoma 2 | mt5 | p.D15N | c.43 G>A | 1 | 1 | Substitution-missense |
| Metastatic histiocytic sarcoma 3 | mt12 | p.L151F | c.451 C>T | 3 | 1 | Substitution-missense |
| Metastatic histiocytic sarcoma 4 | mt15 | p.V198E | c.593 T>A | 4 | 1 | Substitution-missense |
| Metastatic histiocytic sarcoma 5 | mt17 | p.V324I | c.970 G>A | 7 | 1 | Substitution-missense |
| Tumors | Mouses *Arh2* mutation | Amino acid mutation | CDS mutation | Exon | Sample number | Mutation type |
| Lymphoma 1 | mt7 | p.G18S | c.52G>A | 1 | 1 | Substitution-missense |
| Lymphoma 2 | mt7 | p.G18S | c.52G>A | 1 | 1 | Substitution-missense |
|  | mt11 | p.K110K | c.330 A>G | 2 | 1 | Synonymous |
| Lymphoma 3 | mt10 | p.A67V | c.200 C>T | 1 | 1 | Substitution-missense |
|  | mt11 | p.K110K | c.330 A>G | 2 | 1 | Synonymous |
| Lymphoma 4 | mt14 | p.A182T | c.544 G>A | 3 | 1 | Substitution-missense |
| Lymphoma 5 | mt16 | p.R207S | c.621 G>C | 4 | 1 | Substitution-missense |
| Lymphoma 6 | mt18 | p.A344T | c.1030 G>A | 7 | 1 | Substitution-missense |
| Metastatic lymphoma 1 | mt7 | p.G18S | c.52G>A | 1 | 1 | Substitution-missense |
| Metastatic lymphoma 4 | mt14 | p.A182T | c.544 G>A | 3 | 1 | Substitution-missense |
| Metastatic lymphoma 5 | mt16 | p.R207S | c.621 G>C | 4 | 1 | Substitution-missense |
| Metastatic lymphoma 6 | mt18 | p.A344T | c.1030 G>A | 7 | 1 | Substitution-missense |
| Tumors | Mouses *Arh2* mutation | Amino acid mutation | CDS mutation | Exon | Sample number | Mutation type |
| Hepatocellular carcinoma 1 | mt6 | p.L17F | c.49 C>T | 1 | 1 | Substitution-missense |
| Hepatocellular carcinoma 2 | mt6 | p.L17F | c.49 C>T | 1 | 1 | Substitution-missense |
| Hepatocellular carcinoma 3 | mt8 | p.R51T | c.152 G>C | 1 | 1 | Substitution-missense |
|  | mt9 | p.T58T | c.174 C>T | 1 | 1 | Synonymous |
| Hepatocellular carcinoma 4 | mt8 | p.R51T | c.152 G>C | 1 | 1 | Substitution-missense |
|  | mt9 | p.T58T | c.174 C>T | 1 | 1 | Synonymous |
| Hepatocellular carcinoma 5 | mt13 | p.T162I | c.485 C>T | 3 | 1 | Substitution-missense |

**Supplemental Table 3.** Same amino acid mutations sites between human *ARH2* mutations in cancers and murine *Arh2* mutations derived from heterozygous tumors and tumors in nude mice injected with *Arh2*-Het MEFs.

| AA Position | CDS Mutation | *AA Mutation | Legacy Mutation ID | Mutation Type | Count | Primary tissue | Histology | Tumor in *Arh2* Het<sup>#2</sup> |
| --- | --- | --- | --- | --- | --- | --- | --- | --- |
| 7 | c.21G>A | p.A7A | COSM4575853 | Synonymous | 1 | Bone | Ewings sarcoma | mt2: Het1 MEFs and tumor in nude mice |
| 15 | c.43G>A | p.D15N | COSM6595356 | Substitution - Missense | 1 | Large intestine | Adenocarcinoma | mt5: histiocytic sarcoma |
| 17 | c.49C>T | p.L17F | COSM8093697 | Substitution - Missense | 1 | Skin | Malignant melanoma | mt6: hepatocellular Cacinoma |
| 18 | c.52G>A | p.G18S | COSM5473447 | Substitution - Missense | 1 | Large intestine | Adenocarcinoma | mt7: lymphoma |
| 59 | c.177C>T | p.T59T | COSM9763101 | Synonymous | 1 | Skin | Basal cell carcinoma | mt9: hepatocellular cacinoma |
| 68 | c.203C>T | p.A68V | COSM1365603 | Substitution - Missense | 1 | Large intestine | Adenocarcinoma | mt10: lymphoma |
| 116 | c.347T>G | p.L116R | COSM4046200 | Substitution - Missense | 1 | Stomach | Adenocarcinoma | mt3: Het1 MEF and tumor in nude mice |
| 152 | c.454C>T | p.L152F | COSM224459 | Substitution - Missense | 2 | Brain | Glioma | mt12: histiocytic sarcoma |
|  |  |  |  | Substitution - Missense | 1 | Skin | Malignant melanoma | mt12: histiocytic sarcoma |
| 163 | c.488C>T | p.T163I | COSM7354619 | Substitution - Missense | 1 | Lung | Squamous cell carcinoma | mt13: hepatocellular cacinoma |
|  |  |  |  | Substitution - Missense | 1 | Lung | Non small cell carcinoma | mt13: hepatocellular cacinoma |
|  |  |  |  | Substitution - Missense | 1 | Lymphoid | Leukaemia | mt13: hepatocellular cacinoma |
| 183 | c.547G>A | p.A183T | COSM4878741 | Substitution - Missense | 2 | Prostate | Adenocarcinoma | mt14: lymphoma |
| 208 | c.623G>T | p.R208M | COSM8321490 | Substitution - Missense | 1 | Upper aerodigestive tract | Squamous cell carcinoma | mt16: lymphoma |
| 325 | c.973G>A | p.V325I | COSM3786143 | Substitution - Missense | 2 | Pancreas | Ductal carcinoma | mt17: histiocytic sarcoma |
|  |  |  |  | Substitution - Missense | 1 | Lymphoid | Leukaemia | mt17: histiocytic sarcoma |
| 344 | c.1030G>A | p.A344T | COSM1235767 | Substitution - Missense | 1 | Large intestine | Adenocarcinoma | mt18: lymphoma |
|  |  |  |  | Substitution - Missense | 1 | Lymphoid | Plasma cell myeloma | mt18: lymphoma |

**Supplemental Table 4.**
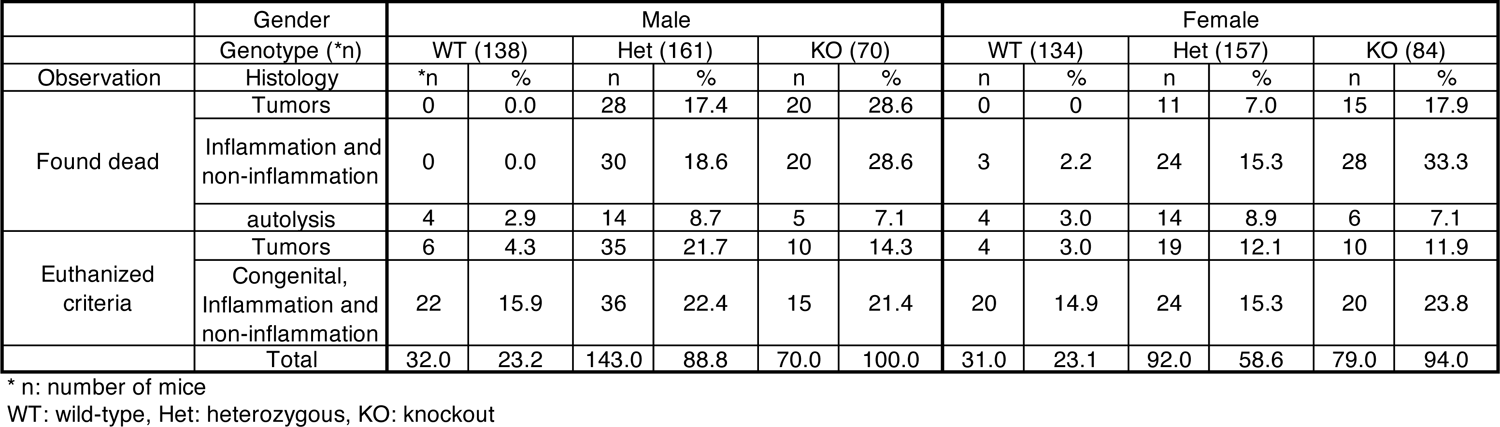
Incidence of unexpected deaths (e.g., tumors, inflammation, non-inflammation, congenital, autolysis) in *Arh2* mice.

**Supplemental Table 5.**
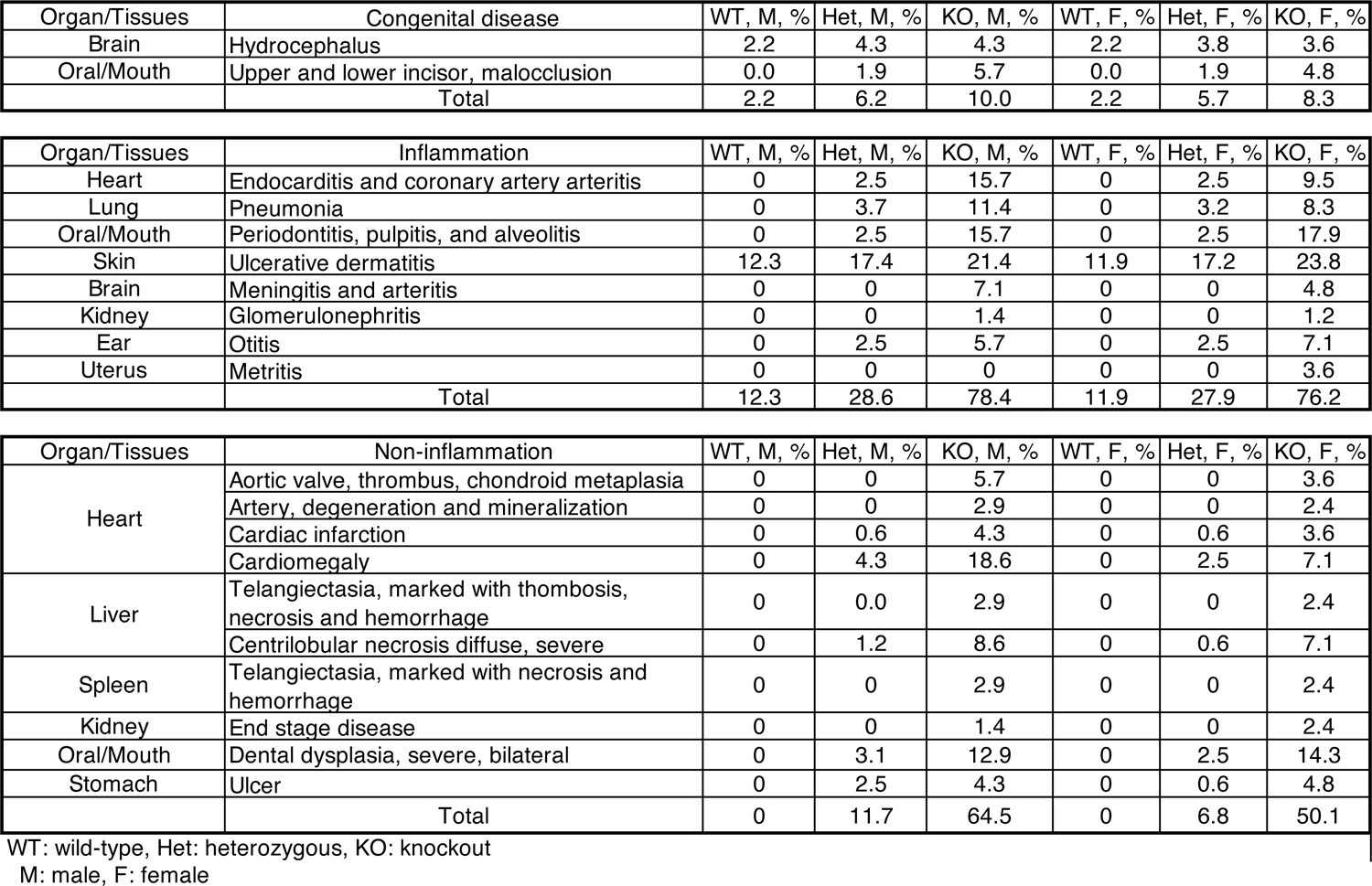
Incidence of unexpected deaths caused by non-tumor diseases (e.g., inflammation, non-inflammation, congenital) in *Arh2* mice.

| Organ/Tissues | Congenital disease | WT, M, % | Het, M, % | KO, M, % | WT, F, % | Het, F, % | KO, F, % |
| --- | --- | --- | --- | --- | --- | --- | --- |
| Brain | Hydrocephalus | 2.2 | 4.3 | 4.3 | 2.2 | 3.8 | 3.6 |
| Oral/Mouth | Upper and lower incisor, malocclusion | 0.0 | 1.9 | 5.7 | 0.0 | 1.9 | 4.8 |
|  | Total | 2.2 | 6.2 | 10.0 | 2.2 | 5.7 | 8.3 |

| Organ/Tissues | Inflammation | WT, M, % | Het, M, % | KO, M, % | WT, F, % | Het, F, % | KO, F, % |
| --- | --- | --- | --- | --- | --- | --- | --- |
| Heart | Endocarditis and coronary artery arteritis | 0 | 2.5 | 15.7 | 0 | 2.5 | 9.5 |
| Lung | Pneumonia | 0 | 3.7 | 11.4 | 0 | 3.2 | 8.3 |
| Oral/Mouth | Periodontitis, pulpitis, and alveolitis | 0 | 2.5 | 15.7 | 0 | 2.5 | 17.9 |
| Skin | Ulcerative dermatitis | 12.3 | 17.4 | 21.4 | 11.9 | 17.2 | 23.8 |
| Brain | Meningitis and arteritis | 0 | 0 | 7.1 | 0 | 0 | 4.8 |
| Kidney | Glomerulonephritis | 0 | 0 | 1.4 | 0 | 0 | 1.2 |
| Ear | Otitis | 0 | 2.5 | 5.7 | 0 | 2.5 | 7.1 |
| Uterus | Metritis | 0 | 0 | 0 | 0 | 0 | 3.6 |
|  | Total | 12.3 | 28.6 | 78.4 | 11.9 | 27.9 | 76.2 |

| Organ/Tissues | Non-inflammation | WT, M, % | Het, M, % | KO, M, % | WT, F, % | Het, F, % | KO, F, % |
| --- | --- | --- | --- | --- | --- | --- | --- |
| Heart | Aortic valve, thrombus, chondroid metaplasia | 0 | 0 | 5.7 | 0 | 0 | 3.6 |
|  | Artery, degeneration and mineralization | 0 | 0 | 2.9 | 0 | 0 | 2.4 |
|  | Cardiac infarction | 0 | 0.6 | 4.3 | 0 | 0.6 | 3.6 |
|  | Cardiomegaly | 0 | 4.3 | 18.6 | 0 | 2.5 | 7.1 |
| Liver | Telangiectasia, marked with thrombosis, necrosis and hemorrhage | 0 | 0.0 | 2.9 | 0 | 0 | 2.4 |
|  | Centrilobular necrosis diffuse, severe | 0 | 1.2 | 8.6 | 0 | 0.6 | 7.1 |
| Spleen | Telangiectasia, marked with necrosis and hemorrhage | 0 | 0 | 2.9 | 0 | 0 | 2.4 |
| Kidney | End stage disease | 0 | 0 | 1.4 | 0 | 0 | 2.4 |
| Oral/Mouth | Dental dysplasia, severe, bilateral | 0 | 3.1 | 12.9 | 0 | 2.5 | 14.3 |
| Stomach | Ulcer | 0 | 2.5 | 4.3 | 0 | 0.6 | 4.8 |
|  | Total | 0 | 11.7 | 64.5 | 0 | 6.8 | 50.1 |

WT: wild-type, Het: heterozygous, KO: knockout M: male, F: female

**Supplemental Table 6.** Incident of mice with tumors in *Arh2* mice

| Gender | Male (%) |  |  | Female (%) |  |  |
| --- | --- | --- | --- | --- | --- | --- |
| Genotype | WT | Het | KO | WT | Het | KO |
| Hepatocellular carcinoma | 0 | 3.1 | 5.7 | 0 | 0.6 | 2.4 |
| Lung adenocarcinoma | 0 | 11.2 | 15.7 | 0 | 3.2 | 7.1 |
| Leukemia | 0 | 0 | 2.9 | 0 | 0 | 1.2 |
| Hemangio sarcoma | 0 | 3.1 | 2.9 | 0 | 1.3 | 2.4 |
| Histiocytic sarcoma | 0 | 12.4 | 11.4 | 0 | 8.9 | 11.9 |
| Lymphoma | 4.3 | 9.3 | 4.3 | 3.0 | 5.1 | 4.8 |
| Total | 4.3 | 39.1 | 42.9 | 3.0 | 19.1 | 29.8 |

**Supplemental Video 1.** *Arh2*-KO mice exhibited decreased myocardial contractility by MRI.

